# A multifunctional polyketide synthase in nematodes produces divergent families of signaling molecules that control different developmental arrests

**DOI:** 10.64898/2026.05.16.725429

**Authors:** Kevin Mai, Chi-Su Yoon, Dilip V. Prajapati, Yuelan Li, Rongrong Yu, Hanh Witte, Subhradeep Bhar, Likui Feng, Elijah Abraham, Matthew T. Gordon, Surasree Rakshit, Priya, Ralf J. Sommer, Rebecca A. Butcher

## Abstract

To improve their chances of reproductive success, nematodes not only must arrest their development in response to adverse growth conditions but also must quickly recover if conditions improve. A polyketide synthase (PKS)-nonribosomal peptide synthetase (NRPS) hybrid assembly line that is expressed in the canal-associated neurons (CANs) of *Caenorhabditis elegans* promotes recovery from starvation-induced larval arrest. Here, we show that in the predatory nematode *Pristionchus pacificus* this assembly line produces a suite of secondary metabolites, including a family of hybrid polyketide-nonribosomal peptides known as the nemamides, the related nemaketides, and a family of ascarylose-modified polyketides named ascarenes. Depending on the starter unit that is loaded onto the PKS, the assembly line can produce dramatically different downstream products. Whereas the nemamides promote recovery from starvation-induced larval arrest, the ascarenes inhibit development of the dauer larval stage and promote recovery. This dichotomy suggests that the PKS-NRPS megasynthetase serves as a signaling hub in the CANs, controlling multiple developmental events. The PKS-NRPS assembly line is highly conserved across many nematode species, and identification of these chemical signals will help to elucidate the signaling pathways that control development in the worm and lead to novel anthelmintics.

## Introduction

Nematodes must assess and integrate multiple factors to determine whether the environment is favorable for development. In response to unfavorable conditions, nematodes can arrest their development at several stages, and if conditions improve, they must rapidly recover to exploit the improved conditions.^1^ The ability to flexibly control development is critical for survival in the boom-or-bust conditions that many nematode species frequently encounter. For example, the free-living bacteriovore *C. elegans* undergoes reproductive growth on rotting plant material, but it can enter the dauer, a stress-resistant dispersal larval stage, if growth conditions become unfavorable.^2^ The chemosensory neurons and signaling pathways that control dauer in *C. elegans* are largely conserved in many parasitic nematode species.^3^ These species have an analogous stage to the dauer, the infective juvenile, which is necessary for host infection.

In *C. elegans*, the dauer is the most thoroughly studied type of larval arrest. If *C. elegans* experiences high population density and low food availability at the second larval (L2) stage, it will not develop into the third larval (L3) stage, but will instead develop into the dauer.^4,5^ If conditions improve, *C. elegans* can recover from the dauer to the fourth larval (L4) stage and resume reproductive growth. In making the dauer decision, *C. elegans* gauges its population density by using chemosensory neurons to detect a secreted pheromone known as the dauer pheromone, which consists of several related ascarosides.^6–10^ The ascarosides are a broad family of glycolipid pheromones that influence not only development, but also variety of behaviors, including mating, aggregation, foraging, and avoidance.^11–13^ The dauer pheromone induces dauer by downregulating the insulin-like/IGF-1 (IIS) pathway and the TGFβ pathway in the worm.^1,14,15^ In addition to the dauer, *C. elegans* can also arrest its development in response to the complete absence of food. If eggs hatch without food, the first larval (L1) stage will arrest its development through a controlled program that is regulated by the IIS pathway.^16,17^ *C. elegans* can also arrest its development at later larval stages (L2-L4) in the complete absence of food.^18,19^

In *C. elegans*, a large hybrid PKS-NRPS megasynthetase, which is encoded by the genes *pks-1* and *nrps-1*, has been implicated in L1 arrest.^20^ The megasynthetase biosynthesizes two hybrid polyketide-nonribosomal peptides known as nemamide A and B in the canal-associated neurons (CANs), two poorly understood neurons that are essential in *C. elegans*.^20–22^ Mutants in *pks-1* and *nrps-1* show reduced survival during L1 arrest and recover more slowing from L1 arrest when returned to food.^20^ These mutants do not up-regulate the expression of specific insulins in arrested L1 larvae in response to food.^20^ However, the effects of the *pks-1* and *nrps-1* mutants are at least partially independent of the DAF-16/FOXO transcription factor, which lies downstream of the insulin/IGF-1 receptor (DAF-2) and plays an essential role in L1 arrest.^20^ The nemamides are endometabolites that are produced at very low levels in the worm,^20^ and hence the mechanism of action of the nemamides has been challenging to study.

The PKS-NRPS megasynthetase that biosynthesizes the nemamides in *C. elegans* is a large, modular enzymatic assembly line.^22,23^ The PKS modules of PKS-1 use acyl transferase (AT) domains to load malonyl subunits onto carrier proteins and ketosynthase (KS) domains to link the subunits together to make the polyketide component of the nemamides. The NRPS modules of NRPS-1 use adenylation (A) domains to load amino acyl subunits onto carrier proteins and condensation (C) domains to link the subunits together to make the nonribosomal peptide component of the nemamides. The presence of a thioesterase (TE) domain at the C-terminus of PKS-1 suggests that biosynthetic intermediates are not directly passed from the upstream PKS-1 to the downstream NRPS-1 but are instead released from PKS-1 before being loaded onto the downstream NRPS-1 to complete nemamide biosynthesis. Many additional accessory enzymes participate in nemamide biosynthesis, including the methyltransferase NEMT-1 and the polyketide-ACP ligase PKAL-1 that traffick intermediates between PKS-1 and NRPS-1.^22^ We hypothesized that this trafficking process may provide the assembly line with biosynthetic flexibility to make additional natural products in addition to the nemamides.

To potentially identify novel PKS-1- and NRPS-1-dependent functions and metabolites, we investigated the satellite model nematode *Pristionchus pacificus*, a nematode species that reproduces on the decaying carcasses of scarab beetles (**Fig. 1a,b**).^24^ Unlike in *C. elegans*, in *P. pacificus*, the CANs are not essential and have been shown to inhibit dauer formation.^25,26^ Using unbiased comparative metabolomics, we show that *pks-1* in *P. pacificus* is required for the production of a suite of different metabolites, including a structurally unique set of nemamides, the structurally related nemaketides, and a family of ascarylose-modified polyketides, the ascarenes. Furthermore, while we show that the nemamides are important for survival during and recovery from starvation-induced larval arrest, we implicate the ascarenes in the control of dauer development. Thus, the PKS-NRPS megasynthetase in *P. pacificus* acts as a signaling hub, biosynthesizing multiple signaling molecules that regulate at least two different developmental arrests.

**Figure 1.**
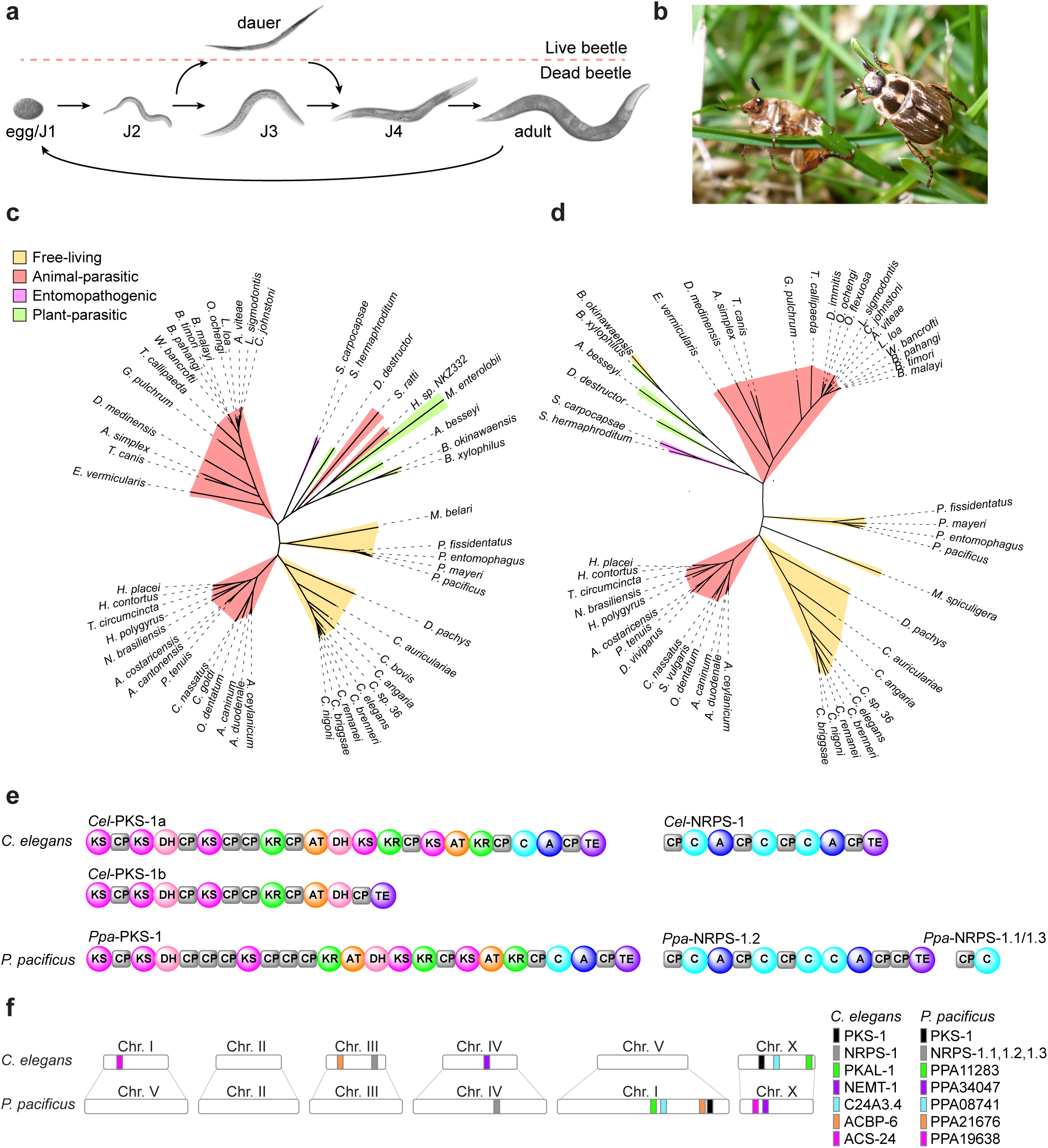
PKS-1 and NRPS-1 homologs in *P. pacificus* and other nematode species. (**a**) Life cycle of *P. pacificus*. The nematode is transported as a dauer on live beetles, and once the beetle dies, the nematode can recover and develop into the reproductive adult. (**b**) The oriental beetle *Exomala orientalist*, a type of scarab beetle, is the primary host of *P. pacificus* in Asia and the United States. (**c,d**) Phylogenetic tree of PKS-1 (**c**) and NRPS-1 (**d**) homologs across different nematode species. The full names of nematode species and protein accession numbers used to generate the PKS-1 homolog phylogenetic tree are in the Methods section. (**e**) Domain organization of PKS-1a, PKS-1b, and NRPS-1 in *C. elegans* and PKS-1 and NPRS-1.1, NRPS-1.2, and NRPS-1.3 in *P. pacificus*. Abbreviations of domains: Carrier protein (CP), acyltransferase (AT), ketosynthase (KS), ketoreductase (KR), dehydratase (DH), adenylation (A), condensation (C), and thioesterase (TE). (**f**) Additional enzymes required for nemamide biosynthesis in *C. elegans* and their chromosomal location, as well as the identities of the closest homologs of these enzymes in *P. pacificus* and their chromosomal location.

## Results

### PKS-1 and NRPS-1 homologs are found in most nematode species

To identify PKS-1 homologs in other nematode species, the *C. elegans* PKS-1 sequence was analyzed by BLAST against nematode genomes in the NCBI database. Only the top BLAST hits from each species were included in the phylogenetic tree, and this tree was filtered by removing hits that were consistent with fatty acid synthase or did not exhibit significant correlation with the domain organization of PKS-1. Homologs of PKS-1 were present in most free-living nematodes, entomopathogenic nematodes, and animal-parasitic nematodes, but were found only in a limited number of plant-parasitic nematodes (**Fig. 1c**). A similar distribution across different nematode species was seen for NRPS-1 homologs (**Fig. 1d**). Thus, it is likely that the ancestral versions of PKS-1 and NRPS-1 were acquired early in nematode evolution.

We hypothesized that in addition to the nemamides, PKS-1 and NRPS-1 might be responsible for the biosynthesis of additional natural products in *C. elegans*. The gene encoding PKS-1, *pks-1*, is predicted to have two splice variants, resulting in longer PKS-1 isoform (PKS-1a), which is responsible for nemamide biosynthesis, and a shorter isoform (PKS-1b), which may produce additional unknown metabolites (**Fig. 1e**). Furthermore, PKS-1a and PKS-1b terminate in a C-terminal TE domain, which likely cleaves intermediates from the assembly line. However, we have been unable to identify additional PKS-1-dependent metabolites in *C. elegans*. Therefore, we decided to explore the natural products produced by PKS-1 homologs in other nematode species, and we focused on *P. pacificus*, a beetle-associated nematode species that has predatory adaptations enabling it to feed not only on the microorganisms on the beetle carcass, but also on other nematodes.^24^

The domain organization of *Ppa*-PKS-1 and *Ppa*-NRPS-1.2 is similar to that of *Cel*-PKS-1a and *Cel*-NRPS-1, respectively (**Fig. 1e**). In addition to *Ppa*-NRPS-1.2, *P. pacificus* contains two truncated NRPS-1 homologs, *Ppa*-NRPS-1.1 and *Ppa*-NRPS-1.3, which are identical to each other and include only a carrier protein and a condensation domain (**Fig. 1e**). Homologs of the many accessory enzymes that are required for nemamide biosynthesis in *C. elegans* are present in *P. pacificus*, but the chromosomal location of the genes that encode them is not conserved, further suggesting that the pathway appeared early in nematode evolution (**Fig. 1f**).

### Identification of nemamides and nemamide-like metabolites in *P. pacificus*

To identify PKS-1-dependent metabolites in *P. pacificus*, we utilized CRISPR-Cas9 gene editing to produce two *pks-1* mutant worm strains, *Ppa*-*pks-1*(*tu1297*) and *Ppa-pks-1*(*tu1298*), both of which contain short deletions near the 5′-end of the gene. Extensive efforts to generate a mutant for *Ppa-nrps-1.2* were unsuccessful. Next, we compared the endometabolomes of wild-type and *pks-1*(*tu1297*) strains via untargeted comparative metabolomics on a LC-high resolution (HR)-MS/MS (**Fig. 2a**). For our metabolomic analysis, we used the commercial software MetaboScape, which merges redundant features that represent different isotopes and adducts of the same metabolite, and we further refined our datasets through manual curation. We first analyzed the metabolites that ionized in positive mode. The volcano plot identified several metabolites that are present in wild type and completely absent in the *pks-1* mutant endometabolome (**Fig. 2b**). Two of the metabolites were named nemamide C and nemamide D, since they both produced similar MS/MS fragmentation patterns to nemamide A and B from *C. elegans*, and another was named nemaketide C, as it is a polyketide that structurally corresponds to the side chain of nemamide C (**Fig. 2c**).

**Figure 2.**
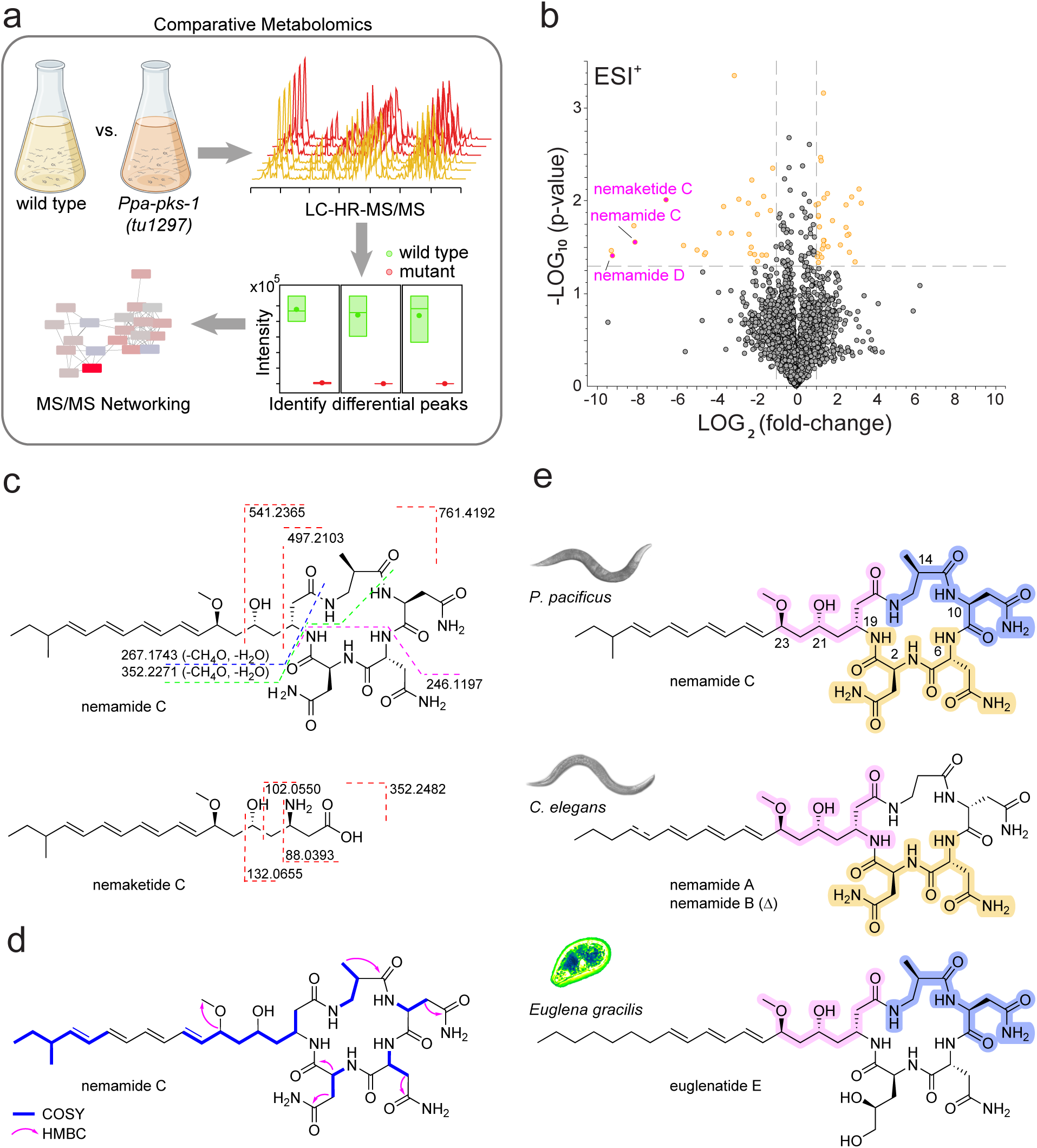
Analysis of PKS-1-dependent metabolites in *P. pacificus* identifies nemamides. (**a**) Strategy for identifying PKS-1-dependent metabolites. Mixed stage cultures of wild-type and *pks-1(tu1297)* worms were grown, the endo- and exometabolites were analyzed by LC-HR-MS/MS, and differential metabolites were identified and characterized using MS/MS networking. (**b**) Volcano plot comparing the average intensities of metabolites in wild-type versus *pks-1*(*tu1297*) endometabolomes in positive (ESI^+^) mode. The x-axis indicates the log ratio of the area of a given peak in wild-type versus *pks-1*(*tu1297*), and the y-axis indicates the statistical significance. (**c**) MS/MS fragmentation patterns of nemamide C and nemaketide C. (**d**) Key NMR correlations from the DQF-COSY and HMBC spectra of nemamide C used in its structure elucidation. (**e**) Assignment of the absolute configuration of nemamide C based in part on the similarity to nemamides A and B from *C. elegans* and euglenatide E and other euglenatides from *E. gracilis*.

In order to determine the chemical structures of the nemamides from *P. pacificus*, we grew mixed staged *P. pacificus* in an axenic, semi-defined medium known as *C. elegans* Habituation and Reproduction (CeHR) medium to generate high-density cultures.^27^ After the extraction of the dried worms, the PKS-1-dependent metabolites were purified using four chromatography steps, silica gel chromatography, HP20ss chromatography, LH-20 chromatography and HPLC. Approximately 50 L of worm culture were processed to purify these compounds in preparation for NMR spectroscopy. Due to the instability of the compounds in the impure form, the worms were processed rapidly in 3L batches through all chromatography steps. Only microgram amounts of nemamide C could be isolated, as it was even less abundant than nemamides A and B from *C. elegans*.

The UV spectrum of nemamide C demonstrated that it, like nemamide B from *C. elegans*, is a tetraene. In comparison to the MS/MS spectrum of nemamide B, the MS/MS spectrum of nemamide C suggests that it has one additional methylene group in its polyketide side chain and one additional methylene group in its macrolactam ring (**Fig. 2c**, **Fig. S1**). The NMR data of nemamide C are highly similar to those of nemamide A and B (**Fig. 2d**, **Fig. S2**, **Table S1**). These data demonstrate that the additional methylene in the polyketide side chain is due to a methyl branch at the ω-2 position and the additional methylene in the macrolactam ring is due to β-aminoisobutyric acid (βAib) replacing the β-alanine that is found in nemamide A and B. Interestingly, a βAib residue is found at this position in the macrolactam ring of the euglenatides, a nemamide-like family of natural products from the algae *Euglena gracilis*.^28^ The similarity of the polyene region of nemamide C with that of nemamide B suggests that the double bonds are all-*trans*. The chemical shifts in nemamide C were compared to those of nemamide A and B, as well as those of the euglenatides from *E. gracilis* (**Fig. 2e**, **Table S2-S3**).^28^ Based on this analysis, the absolute configurations of the stereocenters at positions 2, 6, 19, 21, and 23 of nemamide C likely match those of nemamides A and B (**Fig. 2e, Table S2, S3**). Meanwhile, the absolute configurations of the stereocenters at positions 10 and 14 of nemamide C could not be assigned easily. However, the chemical shifts in this region were more similar to those of the euglenatides, such as euglenatide E, than those of nemamides A and B (**Fig. 2e, Table S2, S3**). Thus, we synthesized four model cyclic peptides, in which the polyketide tail of nemamide C was truncated as a methyl group, and the configurations corresponding to positions 2, 6, and 19 of nemamide C were matched those of nemamide A and B, and the configurations corresponding to positions 10 and 14 were varied (**Fig. S3**). Comparison of the chemical shifts of these four peptides to nemamide C confirmed that the configuration of nemamide C as 2*S*,6*R*,10*S*,14*R*,19*R*,21*R*,23*S* (**Fig. 2e**, **Table S4-S5**).

Analysis of the MS/MS fragmentation patterns of nemaketide C and nemamide D enabled us to propose their chemical structures. The exact mass and MS/MS spectrum of nemaketide C suggest that it corresponds to the side chain of nemamide C (**Fig. 2c, Fig. S1**). Analogous intermediates, referred to here as nemaketides A and B (and previously referred to as nematides A and B^29^), accumulate in the *pkal-1* mutant of *C. elegans*, which is defective for trafficking intermediates between PKS-1 and NRPS-1 in *C. elegans*.^22^ Interestingly, however, whereas nemaketides A and B have not been detected in wild-type *C. elegans*, only in the *pkal-1* mutant, nemaketide C is produced in wild-type *P. pacificus*, indicating that it is not simply a biosynthetic intermediate, but may have an independent signaling function. Comparing the exact mass and MS/MS spectra of nemamide C and nemamide D, we deduced that nemamide D is likely the ring-opened form of nemamide C, but may also have additional structural differences from nemamide C (**Fig. S1**).

### Identification of the ascarenes in *P. pacificus*

In our comparison of the endometabolomes of wild-type and *pks-1*(*tu1297*) strains, we next focused on the metabolites that ionize in negative mode. The volcano plot identified not only the nemamides, but also a different family of metabolites that are present in wild type and completely absent in the *pks-1* mutant endometabolome (**Fig. 3a**). These metabolites, which we named ascarenes A-D, have a different fragmentation pattern from the nemamides and cluster together based on their MS/MS fragmentation patterns (**Fig. 3b**). We again utilized CeHR medium to grow high-density worm cultures for isolation of the ascarenes for structural characterization. The ascarenes are even less abundant than the nemamides, and we only targeted the most abundant of these metabolites, ascarene A, for purification and structure elucidation (**Fig. 3c**). Unfortunately, when *P. pacificus* was propagated over multiple generations in CeHR, the production of ascarene A, unlike that of nemamide C, gradually declined, and we were forced to regularly re-inoculate CeHR with freshly generated *P. pacificus* eggs from bacteria-fed cultures and re-scale up the cultures to obtain batches of worms for extraction.

**Figure 3.**
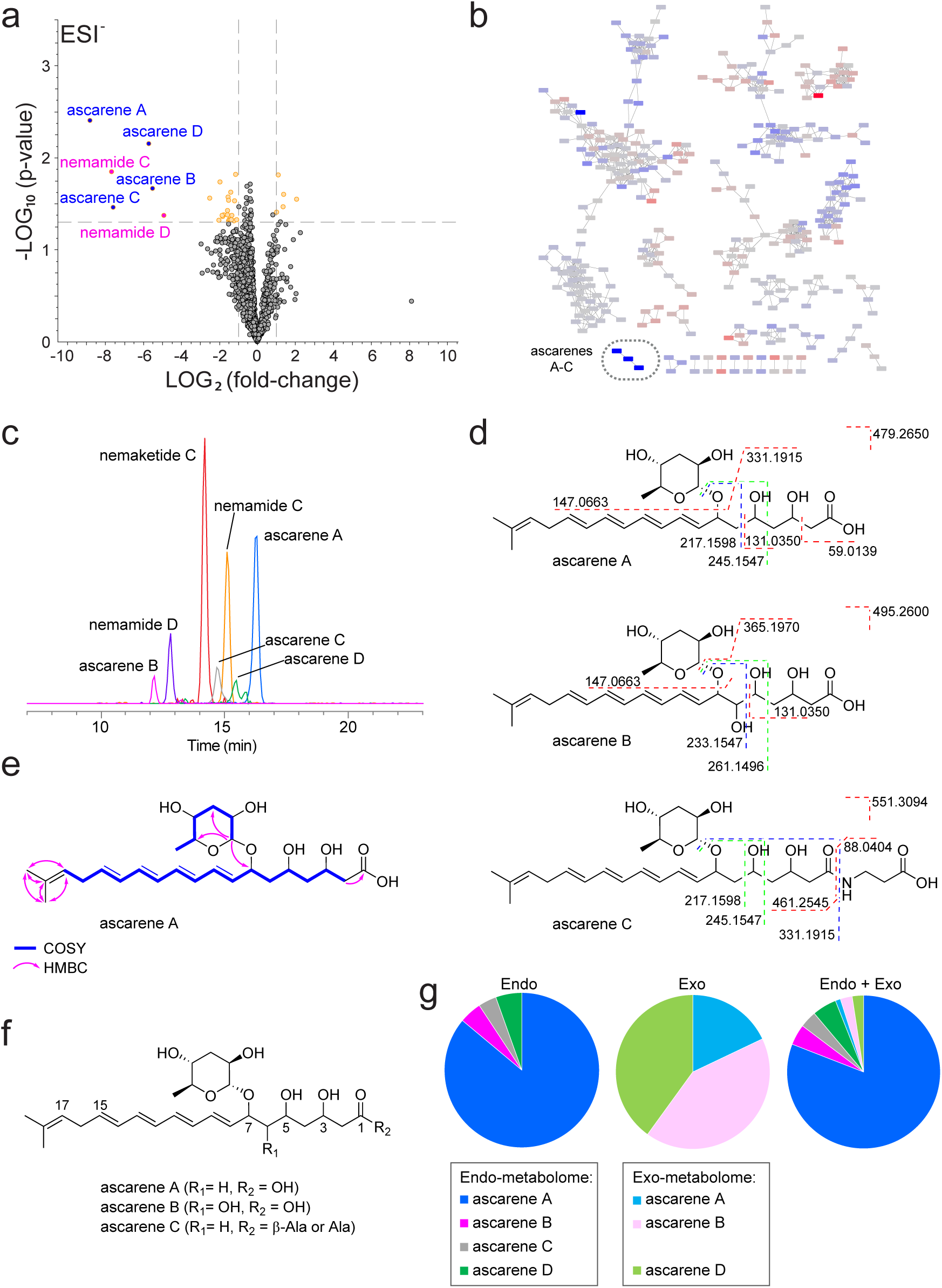
Analysis of PKS-1-dependent metabolites in *P. pacificus* identifies ascarenes. (**a**) Volcano plot comparing the average intensities of metabolites in wild-type versus *pks-1*(*tu1297*) endometabolomes in negative (ESI^-^) mode. The x-axis indicates the log ratio of the area of a given peak in wild-type versus *pks-1*(*tu1297*), and the y-axis indicates the statistical significance. (**b**) Part of the MS/MS network of the data from (**a**). The log ratio of the area of a given peak in wild-type versus *pks-1*(*tu1297*) is indicated using a blue-to-red color scheme, where blue indicates a lower abundance in mutant while red indicates a higher abundance in mutant, relative to wild type. Ascarenes A-C cluster together. Ascarene D had a low intensity and did not have sufficient MS/MS fragmentation to enable clustering. (**c**) Extracted ion chromatograms of nemaketide C (*m/z* 352.2489) in positive (ESI+) mode and nemamide C (*m/z* 759.4068), nemamide D (*m/z* 777.4169), ascarene A (*m/z* 479.2669), ascarene B (*m/z* 495.2615), ascarene C (*m/z* 550.3041), and ascarene D (*m/z* 477.2515) in negative (ESI-) mode. (**f**) MS/MS fragmentation patterns of ascarenes A-C. (**f**) Key NMR correlations from the DQF-COSY and HMBC spectra of ascarene A used in its structure elucidation. (**g**) Chemical structures of ascarenes A-C. (**h**) Relative abundances of the ascarenes in the endometabolome, exometabolome, and endo- and exometabolomes combined.

By monitoring the tetraene UV signal of ascarene A after each column in our purification scheme, we could determine that this compound did not degrade significantly during the purification process. However, once pure and at a comparatively high concentration, the compound rapidly degraded. From a small-scale purification, we were able to isolate a small amount of pure ascarene A that did not degrade, possibly due to its low concentration, and we used this sample to obtain a ^1^H-NMR spectrum in 1.7 mm microcryoprobe (**Fig. S4**). In a subsequent large-scale purification of ascarene A, we elected to collect 2D NMR data, including DQF-COSY, HSQC, HMBC, and NOESY, on semi-purified ascarene A, utilizing our initial ^1^H-NMR spectrum as a guide to focus on the ascarene A-derived NMR correlations (**Fig. S4, Table S6**). Analysis of the MS/MS fragmentation pattern of ascarene A indicated that it consists of a dideoxysugar and a polyketide fragment (**Fig. 3d, Fig. S5**). Based on the NMR data, we were able to determine the structure of the polyketide portion, which has hydroxyl groups at C-3, C-5, and C-7, a tetraene, and a terminal isobutenyl group (**Fig. 3e,f**, **Table S6**). The coupling constants between the protons in the tetraene fragment enabled us to determine the configuration of all of the double bonds, except for the one between C-8 and C-9 due to chemical shift overlap. However, similarities between the chemical shifts of ascarene A and other natural products with an all-*trans* tetraene suggests that the tetraene in ascarene A is all-*trans*. A key HMBC correlation indicates that the polyketide is connected at the C-7 hydroxyl to the anomeric position of a 3,6-dideoxysugar (**Fig. 3e,f**). Through analysis of the NMR data of this sugar, as well as hydrolysis of a small portion of pure ascarene A and sugar derivatization, we were able to determine that this sugar is ascarylose (**Table S7, Fig. S6**).

Similar to ascarene A, ascarenes B and C are tetraenes. Based on the structure of ascarene A, we were able to propose structures for ascarenes B and C utilizing MS/MS data (**Fig. 3d**, **Fig. S5**). The exact mass of ascarene B revealed that it is very similar to ascarene A, but contains an additional oxygen atom. In the MS/MS data of ascarene B, we observed two important MS/MS fragments, *m/z* 261.1483 and *m/z* 285.1867, which suggested the likely location of an additional hydroxyl group (**Fig. 3d**, **Fig. S5**). The exact mass and MS/MS spectrum of ascarene C revealed that it is very similar to ascarene A, except that its carboxylic acid is likely coupled through an amide bond to an alanine/β-alanine amino acid. There are three MS/MS fragments that indicate alanine/β-alanine is present (*m/z* 88.0404) and its likely location (*m/z* 313.1815 and *m/z* 461.2571) (**Fig. 3d**, **Fig. S5**). Ascarene D contains two less hydrogens compared to ascarene A, indicating an additional degree of unsaturation. In the absence of MS/MS data, however, we were unable to propose a structure for ascarene D.

Comparative metabolomics of the exometabolome of wild-type and *pks-1*(*tu1297*) strains revealed that while nemamides C and D and nemaketide C are not found in the exometabolome at all, the ascarenes, are found in the exometabolome to a small degree in mixed-stage cultures (**Fig. S7-S8**). Although present in the exometabolome, the ascarenes are predominantly retained within the worm, with only about 6% found in the exometabolome (**Fig. 3g**). Whereas ascarene A is the predominant ascarene in the endometabolome, ascarene B and D are the predominant ascarenes in the exometabolome. Meanwhile, ascarene C is only found in the endometabolome.

### Divergent biosynthesis of the nemamides/nemaketides and ascarenes

Remarkably, the PKS-NRPS assembly line from *P. pacificus* can biosynthesize two different families of metabolites, the nemamides/nemaketides and the ascarenes. The overall polyketide framework of the two families of metabolites contains some similarities, as highlighted in pink for the nemamides/nemaketides and blue for the ascarenes (**Fig. 4**). The two families differ, however, in their starter unit. Whereas the starter unit for the nemamides/nemaketides likely comes from isoleucine, the starter unit for the ascarenes likely comes from valine. The β,γ-position of the double bond at C17-C18 in the ascarenes (see **Fig. 3f**) suggests that either a β,γ-double bond is directly installed in the growing polyketide by a noncanonical DH shift domain, as in rhizoxin biosynthesis, or a α,β-double bond is first installed and then isomerized to the β,γ-position by a dual function dehydratase/isomerase DH domain, as in gephyronic acid biosynthesis.^30–32^

**Figure 4.**
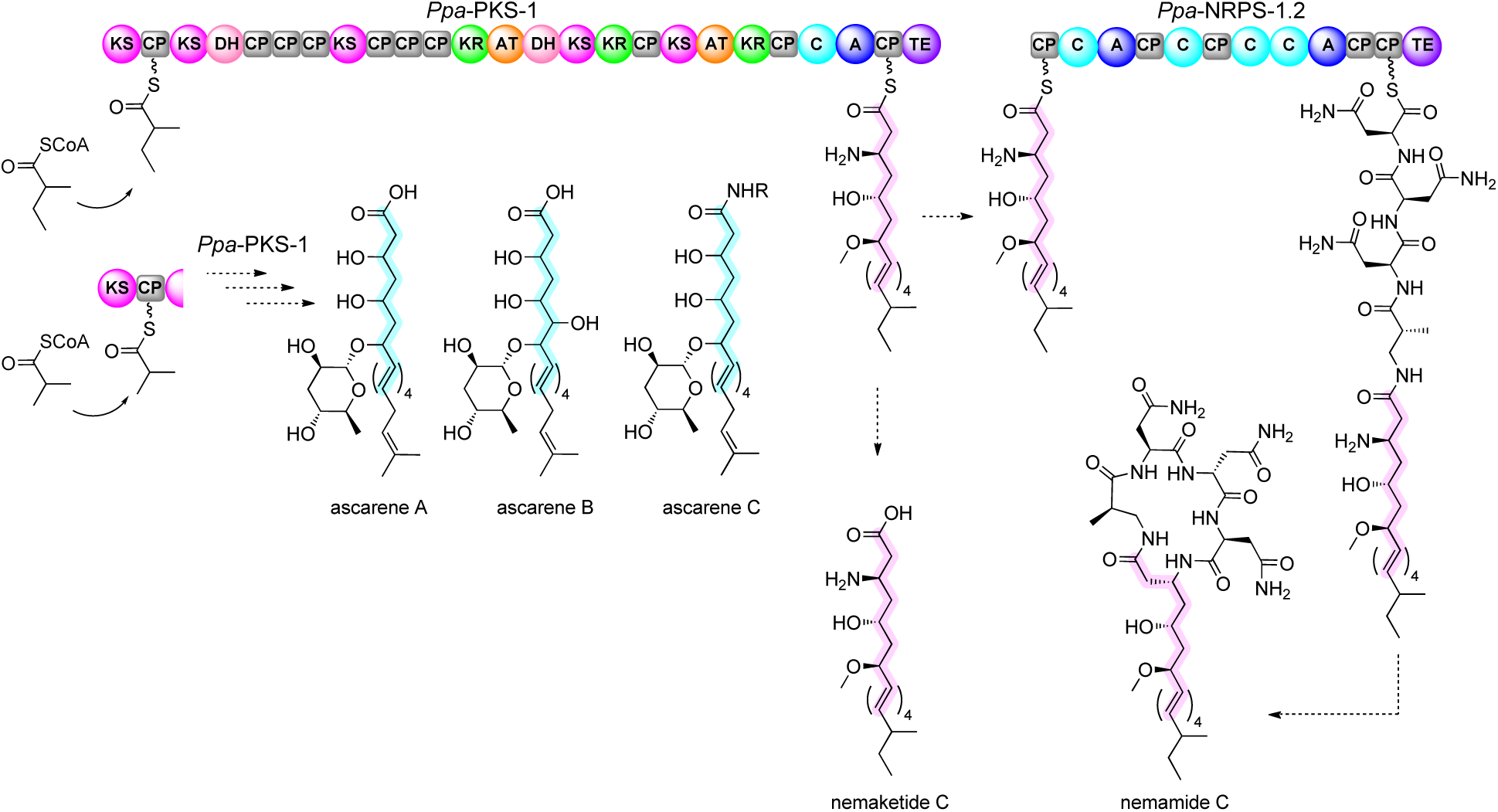
Model for the biosynthesis of the nemamides and ascarenes. The shared chemical framework is highlighted in blue for nemaketide C and nemamide C and in pink for ascarenes A-C. Different starter units are loaded onto PKS-1, leading to either nemaketide C and nemamide C or ascarenes A-C (R = β-Ala or Ala). Nemaketide C produced by PKS-1 can be loaded onto NRPS-1.2, likely by the homolog of PKAL-1 in *P. pacificus*, to produce nemamide C.

The nemamides/nemaketides and ascarenes differ in that the former are modified with a methyl group and the latter are modified with an ascarylose sugar at C7. In *C. elegans*, the methyltransferase that is responsible for methylation in the nemamides/nemaketides has been identified as NEMT-1, but whether methylation occurs during polyketide extension on PKS-1 or during post-synthetic tailoring is not known definitively.^22^ Meanwhile, the ascarenes are likely biosynthesized through modification of the polyketide core with a nucleotide diphosphate (NDP)-activated ascarylose sugar, either during polyketide extension on PKS-1 or during post-synthetic tailoring. The ascarenes have some superficial structural similarity to the ascaroside pheromones.

However, the ascarosides and ascarenes are biosynthesized through completely different pathways. Whereas the ascarenes are biosynthesized through attachment of an ascarylose sugar to a polyketide, the ascarosides are biosynthesized from long-chain ascarosides that have their fatty acid-derived side chains shortened through peroxisomal β-oxidation to generate ascarosides with short- or medium-length side chains.^33–35^ The presence of a C-terminal TE domain on PKS-1 likely allows the release of the nemaketides/ascarenes from PKS-1, as this domain has already been shown to be necessary for the biosynthesis of the nemaketides that accumulate in the *pkal-1* mutant in *C. elegans*.^22^ Thus, this TE domain appears essential for the ability of the PKS-NRPS assembly line to produce multiple families of natural products (nemamides/nemaketides and ascarenes).

### Role of the PKS-NRPS assembly line in starvation-induced larval arrest

Given that the nemamides/nemaketides and ascarenes are produced in such small amounts in the worm and given that they appear to be largely endometabolites, we studied their function primarily though the analysis of mutant worm strains. In addition to the *Ppa*-*pks-1*(*tu1297*) and *Ppa-pks-1*(*tu1298*) mutant strains, also available to us were several previously generated *Cel-pks-1* and *Cel-nrps-1* mutants, including one that should affect both PKS-1a and PKS-1b, *Cel*-*pks-1(ok3769)*, and one that should only affect PKS-1a, *Cel*-*pks-1(ttTi24066)* (**Fig. 5a**).^20^ We first analyzed whether the *Ppa-pks-1* mutant phenotype is similar to that of the *Cel*-*pks-1* and *Cel-nrps-1* mutant phenotypes.

**Figure 5.**
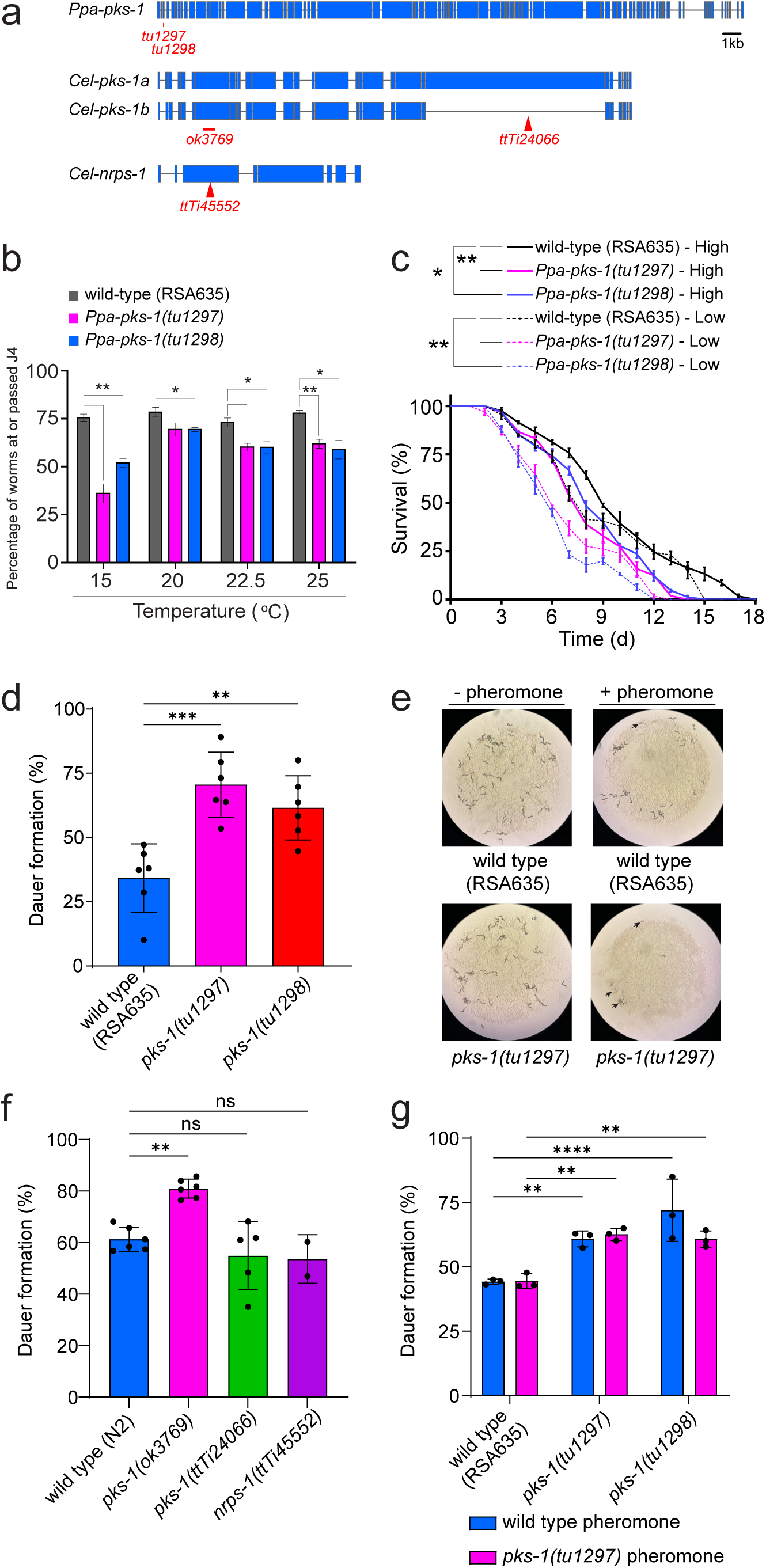
PKS-1-dependent metabolites promote recovery from early larval stage arrest and suppress dauer development. (**a**) Exon structure of *pks-1* from *P. pacificus* and *pks-1* and *nrps-1* from *C. elegans* indicating the location of the mutations in the strains used in this study. (**b**) In *P. pacificus*, percent recovery of wild-type, *pks-1(tu1297)*, and *pks-1(tu1298)* arrested J2 larvae to the J4 stage at different temperatures. (**c**) In *P. pacificus*, survival of wild-type, *pks-1(tu1297)*, and *pks-1(tu1298)* arrested J2 larvae incubated at both low density (0.8-1 worm/μL) and high density (4-6 worms/μL). In (**b**), *p-*values were calculated using an unpaired *t* test (\**p* ≤ 0.05, \*\**p* ≤ 0.01, \*\*\**p* ≤ 0.001, \*\*\*\**p* ≤ 0.0001). In (**c**), the *p*-values were calculated regarding the median lifespan using a method described in the Methods. (**d**) In *P. pacificus*, dauer formation in wild-type, *pks-1(tu1297)*, and *pks-1(tu1298)* worms in response to dauer pheromone generated from wild-type worms. (**e**) Photographs of representative plates from the dauer formation assay in (**d**). The arrows indicate clumps of dauers that are observed in the *pks-1(tu1297)* mutant in the presence of dauer pheromone. (**f**) In *C. elegans*, dauer formation in wild-type, *pks-1(ok3769)*, *pks-1(ttTi24066)*, and *nrps-1(ttTi45552)* mutant worms in response to the synthetic dauer pheromone asc-C6-MK (ascr#2). (**g**) In *P. pacificus*, dauer formation in wild-type and *pks-1(tu1297)* worms in response to dauer pheromone generated from wild-type or *pks-1(tu1297)* worms. Data represent the mean ± standard deviation of three replicates. In (**d**, **f**), *p* values were calculated using one-way ANOVA with Tukey’s multiple comparisons test, and in (**g**), p values were calculated using two-way ANOVA with Dunnett’s multiple comparisons test (\**p* ≤ 0.05, \*\**p* ≤ 0.01, \*\*\**p* ≤ 0.001, \*\*\*\**p* ≤ 0.0001).

*P. pacificus* proceeds through four larval stages (J1-J4) on the way to adulthood, similar to how *C. elegans* proceeds through four larval stages (L1-L4). However, in *P. pacificus*, the J1 stage occurs inside the eggshell and the worms hatch as a J2 (**Fig. 1a**). Thus, to study starvation survival in *P. pacificus*, we hatched the *P. pacificus* eggs in the absence of food and analyzed the survival of the arrested J2 larvae over time. Similar to what we observed in *C. elegans*,^20^ the *Ppa-pks-1* mutant strains of *P. pacificus* were slower to recover from the arrested state (**Fig. 5b**). Also similar to what we observed in *C. elegans*,^20^ the *Ppa-pks-1* mutant strains showed reduced survival during the arrested, starved state compared to wild type, regardless of whether they were maintained at high density or low density, indicating that these effects were likely not due to the absence of a secreted signal but due to the absence of an internal signal (**Fig. 5c**). It is likely that it is the nemamides in *P. pacificus* and *C. elegans* that enhance survival during larval arrest and promote recovery from larval arrest, given that the *Ppa-pks-1* mutants and both *Cel-pks-1* and *Cel-nrps-1* mutants have reduced larval arrest survival and recovery.

### Role of *Ppa*-PKS-1 and *Cel*-PKS-1, but not *Cel*-NRPS-1, in dauer development

In *P. pacificus*, we observed that the *pks-1* mutant worms have enhanced dauer development in the presence of dauer pheromone compared to wild-type (**Fig. 5d,e**). Previously, we reported that in *C. elegans* the *Cel*-*pks-1(ttTi24066)* and *Cel*-*nrps-1(ttTi45552)* mutants displayed a similar amount of dauer formation in response to dauer pheromone as wild type.^20^ This result showed that the nemamides do not affect dauer signaling. Interestingly, however, the *Cel*-*pks-1(ok3769)* mutant is more susceptible to dauer development (**Fig. 5f**). This mutant contains a deletion at the beginning of the *pks-1* gene (presumably preventing the expression of both PKS-1a and PKS-1b) and likely blocks not just the production of the nemamides, but of all natural products produced by *Cel*-PKS-1. Thus, in both *P. pacificus* and *C. elegans* PKS-1 plays a role in the response to dauer pheromone through its role in the production of natural products other than the nemamides. Given that the ascarenes could be produced by the first several domains of PKS-1, the ascarenes are likely candidates for this signal. In *P. pacificus*, mutations in *pks-1* do not affect the potency of crude dauer pheromone (which is generated from the exometabolome of mixed stage cultures) in the dauer formation assay, since the crude dauer pheromone generated from wild-type worms and the crude dauer pheromone generated from *pks-1* mutant worms showed similar abilities to induce dauer in wild-type worms (as well as in *pks-1* mutant worms) (**Fig. 5g**). This result suggests that the *pks-1*-dependent metabolites that suppress dauer formation and/or promote dauer recovery are not found in the exometabolome of mixed stage cultures, and thus they are likely not secreted, but play a signaling role inside the worm.

## Discussion

Here, we have shown that beetle-associated nematode *P. pacificus* uses a PKS-NRPS assembly line to produce a suite of hybrid polyketide-nonribosomal peptide and polyketide natural products, including the nemamides/nemaketides and the ascarenes, and that these natural products promote survival during early larval stage arrest, promote recovery from that arrest, and inhibit dauer development. The similarity of nemamide C from *P. pacificus* to nemamide A/B of *C. elegans* and to the euglenatides of the algae *E. gracilis*, suggest that the nemamides have a fundamental conserved biological function and that nematodes and algae may have acquired the capacity to make these natural products through horizontal gene transfer early in their evolution.^20,28^

The remarkable biosynthetic versatility of PKS-1, which enables it to make dramatically different downstream natural products, likely derives in part from different initiation mechanisms on PKS-1, leading to a downstream bifurcation in the biosynthetic pathway. This bifurcation results in either (1) methylation of the polyketide at the C7 hydroxyl group and installation of the amino group at C3 to produce the nemaketides/nemamides, or (2) modification of the polyketide with an ascarylose sugar at the C7 hydroxyl group and hydroxylation at C3 to produce the ascarenes (**Fig. 4**). PKS-1’s biosynthetic versatility also derives from the unique trafficking mechanism between PKS-1 and the downstream NRPS. We previously hypothesized that in *C. elegans* the C-terminal TE domain of PKS-1 cleaves polyketide products from the assembly line.^22^ This cleavage could lead to the production of polyketides like the nemaketides and ascarenes. Alternatively, reloading of the nemaketides by PKAL-1 onto the N-terminal ACP of NRPS-1 could lead to the production of the nemamides.

Interestingly, we show that in both *P. pacificus* and *C. elegans*, *pks-1* mutants display an enhanced response to dauer pheromone. Since in *C. elegans pks-1* mutants with deletions near the 3′-end of the *pks-1* gene or *nrps-1* mutants do not display an enhanced response to dauer pheromone, it is unlikely that the nemamides play a role in dauer. Furthermore, we have hypothesized that the C-terminal NRPS module of PKS-1 may be involved in installing the amino group in the nemaketides,^22^ and this module would likely be disrupted by *pks-1* mutants with deletions at the 3′-end of the gene (e.g., *pks-1(ttTi24066)*) (**Fig. 5a**). Thus, it is also unlikely that the nemaketides play a role in dauer formation. Since the ascarenes could be biosynthesized by the first several domains of PKS-1, these polyketides may suppress the response to dauer pheromone, possibly by promoting dauer recovery. Unfortunately, the ascarenes are produced in very small amounts in the worm and are highly labile. Our efforts to treat *P. pacificus* worms exogenously with purified ascarene A have not led to reduced dauer formation. Potentially, the ascarenes function only inside in the worm, given that they are for the most part endometabolites and cannot reach their site of action when applied exogenously. Alternatively, multiple ascarenes may need to work synergistically together to affect dauer.

The CANs were previously shown to suppress dauer development in *P. pacificus*, but our results are the first to link the CANs to dauer development in *C. elegans*. In *P. pacificus*, CAN ablation results in increased dauer formation in response to dauer pheromone.^25^ Mutation of the dauerless gene *dau-1*, which is expressed only in the CANs, also results in increased dauer formation.^25^ The dafachronic acids, which oppose dauer formation and promote dauer recovery, are produced in the XXX cells in *C. elegans*, while they are likely produced in the CANs in *P. pacificus*.^26,36^ Thus, there may be differences between the role of the CANs in dauer development in *C. elegans* and *P. pacificus*. However, the role of PKS-1 in these two nematode species appears to be conserved.

Overall, our results suggest that the PKS-NRPS assembly line in the CANs of nematodes acts as a signaling hub and has the biosynthetic potential to produce a range of signaling molecules that impact different aspects of the nematode life cycle. Future studies to understand how these different signaling molecules are produced on the same assembly line will uncover how this hub is regulated. Given the high degree of conservation of the PKS-NRPS assembly line throughout nematode evolution, these studies are likely to further our understanding of the life cycles of both free-living and parasitic nematode species.

## Materials and Methods

### Nematode and bacterial strains

*P. pacificus* strains used in this study were RSA635, RS3471 *pks-1*(*tu1297*), RS3474 *pks-1*(*tu1298*). RS3471 *pks-1*(*tu1297*) and RS3474 *pks-1*(*tu1298*) were generated by CRISPR/Cas9 in the RSA635 background. The *C. elegans* strains used include the wild type N2, as well as RAB9 *pks-1*(*ok3769*), RAB7 *pks-1*(*ttTi24066*), and RAB8 *nrps-1*(*ttTi45552*).^20^ *P. pacificus* and *C. elegans* were grown on nematode growth medium (NGM) agar plates seeded with the *E. coli* strain OP50 or HB101.

### CRISPR/Cas9 generation of *pks-1* mutant strains

CRISPR/Cas9 knockouts were generated using previously published protocols for *P. pacificus*.^37,38^ In brief, crRNAs and tracrRNA (Cat# 1072534) were synthesized by Integrated DNA Technologies (IDT). The CRISPR/Cas9 complex was prepared by mixing 0.5 mg/mL Cas9 nuclease, 0.1 mg/mL tracrRNA, and 0.056 mg/mL crRNA in the TE buffer followed by a 10 min incubation at 37 °C. The *Ppa-eft-3p*::RFP plasmid (20 ng/μL) was added as co-injection marker. The specific sgRNA for knocking out *Ppa-pks-1* is 5′-TGA AGA GCT CGT CGG ATC TG-3′. Microinjections were performed following standard practice using an Eppendorf microinjection system. Injected P0s were removed ca. 16 h post injection. Eggs from each injected P0 animal were allowed to hatch and further transferred onto single NGM agar plates. These F1 animals were genotyped via Sanger sequencing to establish heterozygous null mutant worms. A similar protocol was used to screen homozygous worms in the subsequent F2 generation.

### Phylogenetic analysis

For the phylogenetic trees in **Figure 1c** and **1d**, protein-protein BLAST search of the non-redundant sequence database restricted to Nematoda (taxid:6231) was performed, using *C. elegans* protein sequences as the query.^39^ The sequences were aligned using the default settings of the Clustal Omega multiple sequence alignment tool.^40^ The alignments were subsequently analyzed with the default settings in W-IQ-TREE, a webtool for performing efficient phylogenetic software for Maximum Likelihood (ML) analysis.^41^ The initial parsimony tree was automatically generated by the Phylogenetic Likelihood Library, and a best-fit substitution model was selected via ModelFinder.^42,43^ Evolutionary history was inferred using the ML method and JTT+F+I+G4 substitution model with 1000 ultrafast bootstrap approximations (UFBoot2).^44^ The nearest-neighbor-interchange (NNI) heuristic approach was used to improve the likelihood of each calculated tree. To visualize the results, the phylogenetic tree with the highest log likelihood was uploaded to iTOL.^45^ To identify domains, all sequences were analyzed via the InterProScan webtool using the Pfam database.^46,47^ The resulting domain architecture was visualized using iTOL annotation tools. The resulting PKS-1 trees were pruned to remove BLAST hits that were shorter than 1,000 residues, closely resembled fatty acid synthases, or only contained limited PKS domains. Similarly, the resulting NRPS-1 trees were pruned to remove BLAST hits that did not contain both an A and a C domain. The full names of the nematode species and protein accession numbers used to generate the PKS-1 homolog phylogenetic tree in **Figure 1c**: *Acanthocheilonema viteae* (VBB25232.1), *Ancylostoma caninum* (RCN45771.1), *Ancylostoma ceylanicum* (EYC37446.1), *Ancylostoma duodenale* (KIH69030.1), *Angiostrongylus cantonensis* (KAE9414965.1), *Angiostrongylus costaricensis* (VDM57002.1), *Anisakis simplex* (VDK47117.1), *Aphelenchoides besseyi* (KAI6216298.1), *Brugia malayi* (CRZ22724.1), *Brugia pahangi* (VDN92324.1), *Brugia timori* (VDO06622.1), *Bursaphelenchus okinawaensis* (CAD5229094.1), *Bursaphelenchus xylophilus* (CAD5234868.1), *Caenorhabditis angaria* (CAI5454553.1), *Caenorhabditis auriculariae* (CAD6194341.1), *Caenorhabditis bovis* (CAB3410352.1), *Caenorhabditis brenneri* (EGT30644.1), *Caenorhabditis briggsae* (UMM41728.1), *C. elegans* (NP_508923.2), *Caenorhabditis nigoni* (PIC18413.1), *Caenorhabditis remanei* (KAF1747958.1), *Caenorhabditis sp.* 36 PRJEB53466 (CAI2357108.1), *Cercopithifilaria johnstoni* (CAG9531490.1), *Cylicocyclus nassatus* (CAJ0609431.1), *Cylicostephanus goldi* (VDK40450.1), *Diploscapter pachys* (PAV58557.1), *Ditylenchus destructor* (KAI1718829.1), *Dracunculus medinensis* (VDN58819.1), *Enterobius vermicularis* (VDD86363.1), *Gongylonema pulchrum* (VDN21563.1), *Haemonchus contortus* (CDJ83277.1), *Haemonchus placei* (VDO48929.1), *Halicephalobus sp.* NKZ332 (KAE9548207.1), *Heligmosomoides polygyrus* (VDP18007.1), *Litomosoides sigmodontis* (VDK77668.1), *Loa loa* (EJD75257.1), *Meloidogyne enterolobii* (CAD2171500.1), *Mesorhabditis belari* (CAJ0914828.1), *Nippostrongylus brasiliensis* (WKX99620.1), *Oesophagostomum dentatum* (KHJ99846.1), *Onchocerca ochengi* (VDK64572.1), *Parelaphostrongylus tenuis* (KAJ1359667.1), *Pristionchus entomophagus* (GMT03131.1), *Pristionchus fissidentatus* (GMT33540.1), *Pristionchus mayeri* (GMR33680.1), *Pristionchus pacificus* (KAF8383929.1), *Steinernema carpocapsae* (TKR82333.1), *Steinernema hermaphroditum* (KAK0406688.1), *Strongyloides ratti* (CEF69461.1), *Teladorsagia circumcincta* (PIO69266.1), *Thelazia callipaeda* (VDN04646.1), *Toxocara canis* (VDM41790.1), and *Wuchereria bancrofti* (VDM07833.1). The species names and protein accession numbers used to generate the NRPS-1 homolog phylogenetic tree in **Fig. 1d**: *A. viteae* (VBB26033.1), *A. caninum* (RCN44233.1), *A. ceylanicum* (EPB73156.1), *A. duodenale* (KIH67424.1), *A. costaricensis* (VDM61943.1), *A. simplex* (VDK57631.1), *A. besseyi* (KAI6218735.1), *B. malayi* (VIO98780.1), *B. pahangi* (VDN84132.1), *B. timori* (VDO22169.1), *B. okinawaensis* (CAD5210970.1), *B. xylophilus* (CAD5215369.1), *C. angaria* (CAI5444664.1), *C. auriculariae* (CAD6195629.1), *C. brenneri* (EGT46479.1), *C. briggsae* (CAP32083.2), *C. elegans* (NP_499488.3), *C. nigoni* (PIC42958.1), *C. remanei* (EFP02416.1), *C. sp.* 36 PRJEB53466 (CAI2348997.1), *C. johnstoni* (CAG9535664.1), *C. nassatus* (CAJ0604564.1), *C. goldi* (VDK44017.1), *D. viviparus* (KJH48560.1), *D. pachys* (PAV89777.1), *Dirofilaria immitis* (MCP9258663.1), *D. destructor* (KAI1720343.1), *D. medinensis* (VDN54756.1), *E. vermicularis* (VDD86372.1), *G. pulchrum* (VDK27096.1), *H. contortus* (CDJ82649.1), *H. placei* (VDO04231.1), *H. polygyrus* (VDP54875.1), *L. sigmodontis* (VDK72408.1), *L. loa* (EFO26749.2), *M. spiculigera* (CAJ0579960.1), *N. brasiliensis* (WKY06294.1), *O. dentatum* (KHJ98077.1), *Onchocerca flexuosa* (VDO24867.1), *O. ochengi* (VDK63304.1), *P. tenuis* (KAJ1364105.1), *P. entomophagus* (GMS95933.1), *P. fissidentatus* (GMT24824.1), *P. mayeri* (GMR48516.1), *P. pacificus* (KAF8374438.1), *S. carpocapsae* (TKR83073.1), *S. hermaphroditum* (KAK0394430.1), *Strongylus vulgaris* (VDM67814.1), *T. circumcincta* (PIO73139.1), *T. callipaeda* (VDN03035.1), *T. canis* (VDM39775.1), and *W. bancrofti* (EJW85495.1).

### Extraction of the endometabolome from *P. pacificus*

For small-scale worm extractions, wild-type and *pks-1* mutant worms were grown at room temperature on three NGM agar plates (10 cm) containing 0.75 mL 25X OP50. When the worms were fully saturated in the NGM agar plate, they were transferred to a 2.8 L baffled flask containing 500 mL of S medium. The worm cultures were grown in a shaking incubator at 22.5°C and 200 rpm for 9 d and fed with 10 mL of 25X OP50 daily. The culture flasks were chilled on ice for 30 min to settle the worms. Then, the worms were transferred from the bottom of the flask to a 50 mL centrifuge tube and were centrifuged at 800 g for 5 min to separate the worms from the culture media. Following the separation, the worms were soaked with water for 30 min twice in a shaking incubator at 22.5°C and 200 rpm to remove bacteria from their digestive tract. The washed worms were then frozen for lyophilization. The dried worm pellets were ground with sea sand using a mortar and pestle, and then extracted with 20 mL of 190 proof ethanol for 3 h. The extracts were centrifuged at 800 g for 5 min and the supernatant was dried in vacuo. The endometabolites were resuspended in 120 μL of 50% ethanol/water before LC-MS analysis. The samples were analyzed using a LUNA 5 μm C18 column (100 × 34.6mm; Phenomenex) coupled with an Agilent 6130 single quad mass spectrometer. The following solvent gradient was used with a flow rate of 0.7 mL/min: 95% buffer A, 5% buffer B, 0 min; 0% buffer A, 100% buffer B, 20 min; 0% buffer A, 100% buffer B, 22 min; 95% buffer A, 5% buffer B, 23 min; 95% buffer A, 5% buffer B, 26 min (Buffer A represents water with 0.1% formic acid; Buffer B represents acetonitrile with 0.1% formic acid).

### Extraction of the exometabolome from *P. pacificus*

Wild-type and *pks-1* mutant worms were grown at room temperature on three NGM agar plates (10 cm) containing 0.75 mL 25X OP50. When the worms were fully saturated in the NGM agar plate, they were transferred to a 2.8 L baffled flask containing 500 mL of S medium. The worm cultures were grown in a shaking incubator at 22.5°C and 200 rpm for 9 d and fed with 10 mL of 25X OP50 daily. The culture flasks were chilled on ice for 30 min to settle the worms. Once the worms settled, the exometabolome (secreted metabolites) was collected and centrifuged at 2,675 g for 10 min to remove excess bacterial residue. The bacterial free exometabolome was frozen down in preparation for lyophilization. The dried exometabolome was extracted with methanol and dried down in vacuo for preparation of mass spectrometry analysis. The exometabolites were resuspended in 120 μL 50% methanol/water before LC-MS analysis.

### High-resolution mass spectrometry analysis

All high-resolution LC-MS/MS analysis was performed on a Bruker Impact II QTOF mass spectrometer coupled with an UltiMate 3000 RSLC nano System using a previously described method with the following chromatographic changes^48^. For all experiments on the endometabolome, separation was performed on a capillary column ThermoScientific Acclaim PepMap RSLC, C18 (300 μm x 15 cm, 2 μm, 100 Å) with a C18 precolumn (3 mm x 2 cm,75 μm,100 Å) column, maintained at 30 °C. 10 mM ammonium formate with 0.1% formic acid was mobile phase A, and acetonitrile with 0.1% formic acid was mobile phase B. The following solvent gradient was used, with a flow rate of 5 µL/min: 98% buffer A, 2% buffer B, at 0 min; 25% buffer A, 75% buffer B, at 30 min; 2% buffer A, 98% buffer B, at 35 min and then held constant for next 5 min. For experiments on the exometabolome, separation was performed on a Hypersil GOLD aQ Polar Endcapped C18 3 μm 175Å (2.1 x 150 mm) column, maintained at 30 °C, using 10 mM ammonium formate with 0.1% formic acid as mobile phase A and acetonitrile with 0.1% formic acid as mobile phase B. The following solvent gradient was used with a flow rate of 0.2 mL/min: 95% buffer A, 5% buffer B, at 0 min; 40% buffer A, 60% buffer B, at 20 min; then held constant for the next 7 min. All the samples were analyzed in positive and negative ESI mode. For MS/MS analysis, we acquired all the data in data dependent acquisition (DDA) mode where the mass spectrometer selected the most intense precursor ions at the first stage for further fragmentations. For analysis of the endometabolome, the mass spectrometer parameters were set as follows: end plate offset 500 V, capillary voltage 2.5 kV, dry temperature 200°C, collision RF 300 Vpp, transfer time 40.0 μs, prepulse storage 6μs, collision energy 25.0 eV, and mass range *m/z* 50-1500. For analysis of the exometabolome, the mass spectrometer parameters were set as follows: end plate offset 500 V, capillary voltage 2.5 kV, dry temperature 200°C, collision RF 300 Vpp, transfer time 50.0 μs, prepulse storage 5μs, collision energy 25.0 eV and mass range *m/z* 50-1300.

For all data processing steps and feature detection, MetaboScape (v5.0.0) software platform (Bruker) was used. To extract differential metabolic features from datasets, the raw data files of the biological triplicate samples from the control and mutant group were uploaded onto MetaboScape, and the peaks were selected with a signal intensity greater than 1 x 10^3^ counts and a mass range *m/z* 150-1500. Features with multiple adduct ions were combined, and a t-test was performed to generate a volcano plot to identify differential features. To generate molecular networks, the analyzed feature list from each dataset was exported from MetaboScape as a Feature Based Molecular Networking-GNPS compatible output file and uploaded to the GNPS web server.^49,50^ The following parameters were used to build all the networks: precursor ion mass tolerance: 0.02, fragment ion mass tolerance: 0.02, min pair cosine score: 0.5, minimum fragment ions: 3, maximum connected component size: 100, library search min matched peaks: 4. The generated networks were visualized in Cytoscape (v 3.7.1), and color coded based on the fold change using a log2 scale.

### Purification and characterization of PKS-1-dependent metabolites

To begin purifying and characterizing PKS-1-dependent metabolites, eggs were isolated from gravid young adult worms using the modified alkaline bleach method (10% v/v bleach and 10 N NaOH) and used to inoculate 5 mL of CeHR medium. The culture was grown for 7-10 d in a shaker before transferring the culture to a 250 mL Erlenmeyer flask containing 50 mL of CeHR followed by additional 7-10 d of culturing. This 50 mL culture was transferred to 500 mL of CeHR medium in a 2.8 L baffled flask, and grown for an additional 7-10 d. The CeHR medium contained 20% cow’s milk and cultures were grown in a shaking incubator at 22.5°C and 200 rpm. Worms were then collected by centrifugation at 800 g for 5 min and washed with water twice for 30 min. The worms were frozen at −80°C and stored at that temperature until it was time for lyophilization. After freeze-drying, the worms were pulverized for 10 min with sand (twice the amount of dried worms) using a mortar and pestle. The pulverized worm/sand combination was then extracted with 190 proof ethanol (20 mL per L of worm culture) in an Erlenmeyer flask and shaken at room temperature at 200 rpm for 3 h. The mixture was centrifuged at 800 g for 5 min and the supernatant was filtered through a cotton plugged glass Pasteur pipette. The filtered supernatant was concentrated in vacuo before being subjected to approximately 60 g silica gel for chromatography. The extract was eluted with a gradient of hexane, ethyl acetate, ethyl acetate/methanol (1:1), and methanol (250 mL each) to give four fractions (A-D). Fraction C and D were processed individually, first dried in vacuo, followed by resuspension in 24 mL methanol and centrifuged at 3500 rpm for 10 min. The supernatant was dried and dissolved in 10 mL of 70% methanol/water. The resuspended extract was then applied to a HP-20ss column, eluting with methanol/water (7:3 to 10:0) to give eight subfractions (200 mL each). Subfraction C-3 and D-5 was applied to a Sephadex LH-20 column individually, eluting with methanol to give 20 subfractions containing 18 mL (C3a-C3t, D5a-D5t respectively). Each fraction was dried in vacuo and analyzed using LC-MS to identify the masses of peaks with tetraene UV signatures. Fractions that contained the same compound were combined. The fractions that contained nemamide C were further purified by two steps of HPLC. The first step used a gradient of methanol and water (ramping from 10% to 100% methanol over 30 min, holding 100% methanol for 6 min, then returning to 10% methanol over 4 min with the column (Agilent, eclipse XDB-C18 column, 150 x 4.6mm, 5 μm); flow rate 0.7 mL/min; UV detection at 320 nm. After collect nemamide C from first step, it was further purified using second step used a gradient of acetonitrile and water (ramping from 5% to 100% acetonitrile over 30 min, holding 100% acetonitrile for 5 min, then returning to 5% acetonitrile over 4 min with the column (GL Science, InertSustain-C18 column, 150 x 4.6mm, 5 μm); flow rate 1 mL/min; UV detection at 320 nm. The fractions that contained ascarene A were further purified by one round of HPLC to yield the semi-purified sample for full NMR characterization and an additional round of HPLC to yield the fully purified sample for acid hydrolysis and determination of the absolute configuration of the sugar in the molecule. The first HPLC purification used a gradient of acetonitrile and water (ramping from 15% to 80% acetonitrile over 30 min, from 80% to 100% acetonitrile over 2 min, holding 100% acetonitrile for 4 min, then returning to 15% acetonitrile over 2 min and holding at 15% for 7 min with the column (GL Science, InertSustain-C18 column, 150 x 4.6mm, 5 μm); flow rate 1 mL/min; UV detection at 320 nm), to obtain semi-purified ascarene A for the purposes of 2D NMR analysis. The second HPLC purification was performed using the same method and column, but used methanol and water, to obtain fully purified ascarene A for analysis of the sugar composition. Nemamide C: For NMR spectra and data, see **Figure S2** and **Table S1**; HR-ESIMS (*m/z*): [M-H]^-^ calcd. for C_36_H_55_N_8_O_10_ 759.4041 found 759.4067. Nemamide D: HR-ESIMS (*m/z*): [M-H]^-^ calcd. for C_36_H_57_N_8_O_11_ 777.4147 found 777.4165. Nemaketide C: HR-ESIMS (*m/z*): [M+H]^+^ calcd. for C_20_H_34_NO_4_ 352.2482 found 352.2488. Ascarene A: For NMR spectra and data, see **Figure S4** and **Table S6**; HR-ESIMS (*m/z*): [M-H]^-^ calcd. for C_26_H_39_O_8_ 479.2650 found 479.2669. Ascarene B: HR-ESIMS (*m/z*): [M-H]^-^ calcd. for C_26_H_39_O_9_ 495.2600 found 495.2615. Ascarene C: HR-ESIMS (*m/z*): [M-H]^-^ calcd. for C_29_H_44_NO_9_ 550.3022 found 550.3041. Ascarene D: HR-ESIMS (*m/z*): [M-H]^-^ calcd. for C_26_H_37_O_8_ 477.2494 found 477.2493.

### Model cyclic peptide synthesis

Cyclic peptides **S1-S4** were synthesized on a 0.1 mmol scale using a Liberty Blue automated microwave peptide synthesizer, starting with H-L-Asn(Trt)-2-Cl resin. Cyclic peptide **S1** was synthesized by sequentially coupling H-L-Asn(Trt)-2-Cl resin, Fmoc-D-Asn(Trt)-OH, Fmoc-D-Asn(Trt)-OH, Fmoc-D-βAib-OH, and Fmoc-D-β-HAla-OH; Cyclic peptide **S2** was synthesized by sequentially coupling H-L-Asn(Trt)-2-Cl resin, Fmoc-D-Asn(Trt)-OH, Fmoc-D-Asn(Trt)-OH, Fmoc-L-βAib-OH, and Fmoc-D-β-HAla-OH; Cyclic peptide **S3** was synthesized by sequentially coupling H-L-Asn(Trt)-2-Cl resin, Fmoc-D-Asn(Trt)-OH, Fmoc-L-Asn(Trt)-OH, Fmoc-D-βAib-OH, and Fmoc-D-β-HAla-OH; Cyclic peptide **S4** was synthesized by sequentially coupling H-L-Asn(Trt)-2-Cl resin, Fmoc-D-Asn(Trt)-OH, Fmoc-L-Asn(Trt)-OH, Fmoc-L-βAib-OH, and Fmoc-D-β-HAla-OH. Fmoc-protected amino acids, *N,N′*-diisopropylcarbodiimide (DIC) activator, OxymaPure activator base, and 20% piperidine deprotection solution in N,N-dimethylformamide (DMF) were used for the synthesis. Standard protocols for the synthesizer (resin swelling, coupling cycles, and final deprotection cycles) were used. After the peptide synthesis, the resin was transferred to a glass vessel using 1:1 (v/v) of dichloromethane (DCM) and DMF, rinsed with 4×10 mL DCM and dried. Side-chain cleavage from the resin was achieved by treating with a 1:4 (v/v) of 1,1,1,3,3,3-hexafluoro-2-propanol and DCM for 30 minutes at room temperature. This process was repeated two additional times, and the combined cleavage solution was concentrated under vacuum. For cyclization, the crude linear peptide (0.1 mmol) was dissolved in 60 mL DMF, followed by the addition of 4-(4,6-dimethoxy-1,3,5-triazin-2-yl)-4-methylmorpholinium tetrafluoroborate (DMTMM^+^BF_4_^-^, 66 mg, 0.2 mmol) and *^i^*Pr_2_NEt (26 mg, 0.2 mmol). The solution was stirred overnight and then concentrated under reduced pressure. For deprotection of the trityl groups, 10 mL of a TFA, triisopropylsiline, and water solution (95:2.5:2.5, v/v/v) was added and stirred at room temperature for 2 h, and then concentrated to afford the crude cyclic peptide. The crude cyclic peptide was initially purified on a C18 column (50 g Octadecyl-functionalized silica gel, 3.5 cm × 50 cm), eluted with 10% methanol, and then further purified using reversed-phase HPLC (0–5 min: 2% acetonitrile, 5–15 min: 2–30% acetonitrile, 15–20 min: 30% acetonitrile) to give pure cyclic peptide.

### Acid hydrolysis of ascarene A and determination of the sugar absolute configuration

Ascarene A (one tenth of final purified sample) was hydrolyzed with 0.4 mL of 3 N HCl at 80 ℃ for 2.5 h to obtain the sugar moiety. The hydrolysate was dried using a rotavap and then dissolved in 50 μL of pyridine. To this sample, 150 μL of pyridine containing 150 μg of L-cysteine methyl ester hydrochloride was added, and the mixture was heated to 60 ℃ for 1 h. Subsequently, 0.5 μL of o-tolyl-isothiocyanate was added, and the solution was heated again at 60 ℃ for 1 h. Ascarylose and tyvelose standards (150 μg each) were subjected to the same procedure as the hydrolysate. The mixture was then evaporated and dissolved in methanol for LC-MS analysis. The samples were analyzed using analytical HPLC with a Phenomenex C18 column (5 µm, 4.6 × 100 mm) at a flow rate of 0.7 mL per min. Gradient elution was performed from 5% acetonitrile in water with 0.1% formic acid to 100% acetonitrile with 0.1% formic acid over 20 min. The retention time of sugar portion of ascarene A was 11.91 min, which was the same as that of ascarylose (11.92 min).

### Preparation of dauer pheromone and dauer formation assay

To prepare dauer pheromone, saturated worms from 10 cm plates were transferred into 500 mL of S medium and grown for 9 d, while being fed with 10 mL of 25X OP50 daily. The supernatant was separated from the worms by 800 g centrifugation and followed centrifugation at 2,675 g to remove any residual OP50. The supernatant was stirred with 30 g of activated charcoal overnight. The charcoal was collected and washed several times with water followed by an ethanol extraction. The ethanol extract was concentrated using rotary evaporator and resuspended in 3 mL of 50% ethanol/water. The concentrated dauer pheromone was added in the warm NGM agar, which contained no peptone and cholesterol but included with 50 μg/mL kanamycin (final concentration). Subsequently, 20 μM kanamycin-inactivated OP50 (8 mg/mL) was added to the 3.5 cm NGM agar plates, and young adult *P. pacificus* was placed on the agar plate for 2 h to acquire 60-90 eggs. The plates were incubated for 2 d at 25°C before scoring dauer versus non-dauer. For all the experiments, three biological replicates were performed on the same day. We discovered a consistent pattern within the experiments, which were seen in independent biological replicates on different days.

### J2 starvation survival assay

For starvation survival assays, eggs were isolated from gravid young adult worms using the modified alkaline bleach (10% *v*/*v* of bleach and 10 N NaOH) method previously described. (Werner et al, 2017). The eggs were resuspended with M9 buffer dilution in a 50 mL conical tube and shaken for 24 h at 22.5°C and 200 rpm. Subsequently, the density of the synchronized J2s was adjusted to 4-6 J2s/μL (high density) or 0.8-1 J2s/μL (low density) in M9 buffer. Every day, three 20 μL aliquots of starved J2 samples were taken and seeded on to 3.5 cm NGM plates containing OP50 at room temperature. In 24 h, the plated worms were scanned for carcasses, since it is easier to score. After 2-3 d, the worms that passed the J3 stage were scored as surviving worms. For each experiment, three replicates were used to analyze the average of surviving worm strains daily.

### J2 recovery assay

For the J2 recovery assays, eggs were harvested via the modified alkaline bleach method from gravid adult worms. The harvested eggs were diluted in M9 buffer to give a concentration of 1 egg/μL. The eggs were allowed to hatch in M9 buffer at 22.5°C and 200 rpm in a shaking incubator, where they were maintained for 48 h such that they arrested their development. After 48 h, worms were placed on 3.5 cm NGM agar plates seeded with OP50 and placed at different temperatures (15°C, 20°C, 22.5°C and 25°C). The worms were scored at different time points depending on the incubating temperature (75 h at 15°C, 40 h at 20°C, 34 h at 22.5°C, and 27 h at 25°C). They were then scored as recovered if they were able to reach J4 or higher development in the aliquoted time. Each data set was performed in triplicate.

### Statistical analysis of survival assays

Survival curves were fit with a sigmoidal curve of variable slope using nonlinear regression in GraphPad Prism, setting the maximum survival to 100% and the minimum survival to 0%. The mean survival times were compared using an extra sum-of-squares F test, and the *p*-values reported reflect the likelihood these survival times are different for the compared conditions.

## Acknowledgements

The authors would like to thank Jim Rocca for his help and advice obtaining NMR spectra. The authors would like to thank Matthias Herrman for the beetle photo. This work was supported by grants from the National Institutes of Health (R35 GM144076 (to R.A.B.), the National Science Foundation (CHE-1555050 to R.A.B.), and the Max Planck Society (to R.J.S.).

## Contributions

D.V.P., S.B., and P. performed metabolomics; K.M., C.-S.Y., and L.F. isolated compounds; C.-S.Y., D.V.P., L.F., and Y.L. elucidated the structures of compounds; K.M., Y.L., and S.R. performed biological assays; H.W. generated mutant strains; R.Y. synthesized compounds; R.J.S. and R.A.B. conceived and supervised the study; R.A.B. wrote the manuscript with approval from all authors.

## Notes

### Competing Interest Statement

The authors have declared no competing interest.

### Summary of Updates

Nematides have been renamed nemaketides to help to differentiate them from nemamides.

