## Supplementary Information for "A multifunctional polyketide synthase in nematodes produces divergent families of signaling molecules that control different developmental arrests"

### Supplementary Results

### Table of Contents

**Contents** **Page**


| | | |
|---|---|---|
| Figure S1 | MS/MS spectra of the nemamides of *P. pacificus* | 4 |
| Figure S2 | NMR spectra of nemamide C in dimethyl sulfoxide-$d_6$ | 5 |
| Table S1 | $^1$H and $^{13}$C NMR data derived from $^1$H, dqf-COSY, HSQC, and HMBC spectra for nemamide C in dimethyl sulfoxide-$d_6$ | 11 |
| Table S2 | Comparison of $^1$H NMR chemical shifts of the polyketide chain of nemamide C with the polyketide chains of nemamide A and of euglenatides D and E | 12 |
| Table S3 | Comparison of $^1$H NMR chemical shifts of the peptide head of nemamide C with the peptide head of nemamide A and euglenatides B, C, and E | 13 |
| Figure S3 | Structures of the four model cyclic peptides | 14 |
| Table S4 | $^1$H and $^{13}$C NMR chemical shifts derived from $^1$H, DQF-COSY, HSQC, and HMBC spectra for the four cyclic peptides in dimethyl sulfoxide-$d_6$ | 15 |
| Table S5 | Differences between the cyclic peptides and nemamide C in terms of $^1$H and $^{13}$C NMR chemical shifts | 16 |
| Figure S4 | NMR spectra of ascarene A in methanol-$d_4$ | 17 |
| Table S6 | $^1$H and $^{13}$C NMR data derived from $^1$H, dqf-COSY, HSQC, and HMBC spectra for ascarene A in methanol-$d_4$ | 24 |
| Figure S5 | MS/MS spectra of the ascarenes of *P. pacificus* | 25 |
| Table S7 | Comparison of $^1$H and $^{13}$C NMR chemical shifts of the sugar of ascarene A with ascarylose of asc-C6-MK and paratose of part#9 in methanol-$d_4$ | 27 |
| Figure S6 | Determination of absolute configuration of the sugar in ascarene A | 28 |
| Figure S7 | Comparative metabolomics between wild-type and *pks-1(tu1297)* exometabolomes | 29 |
| Figure S8 | Production of nemamides and ascarenes in the endometabolome and exometabolome of *P. pacificus* | 30 |

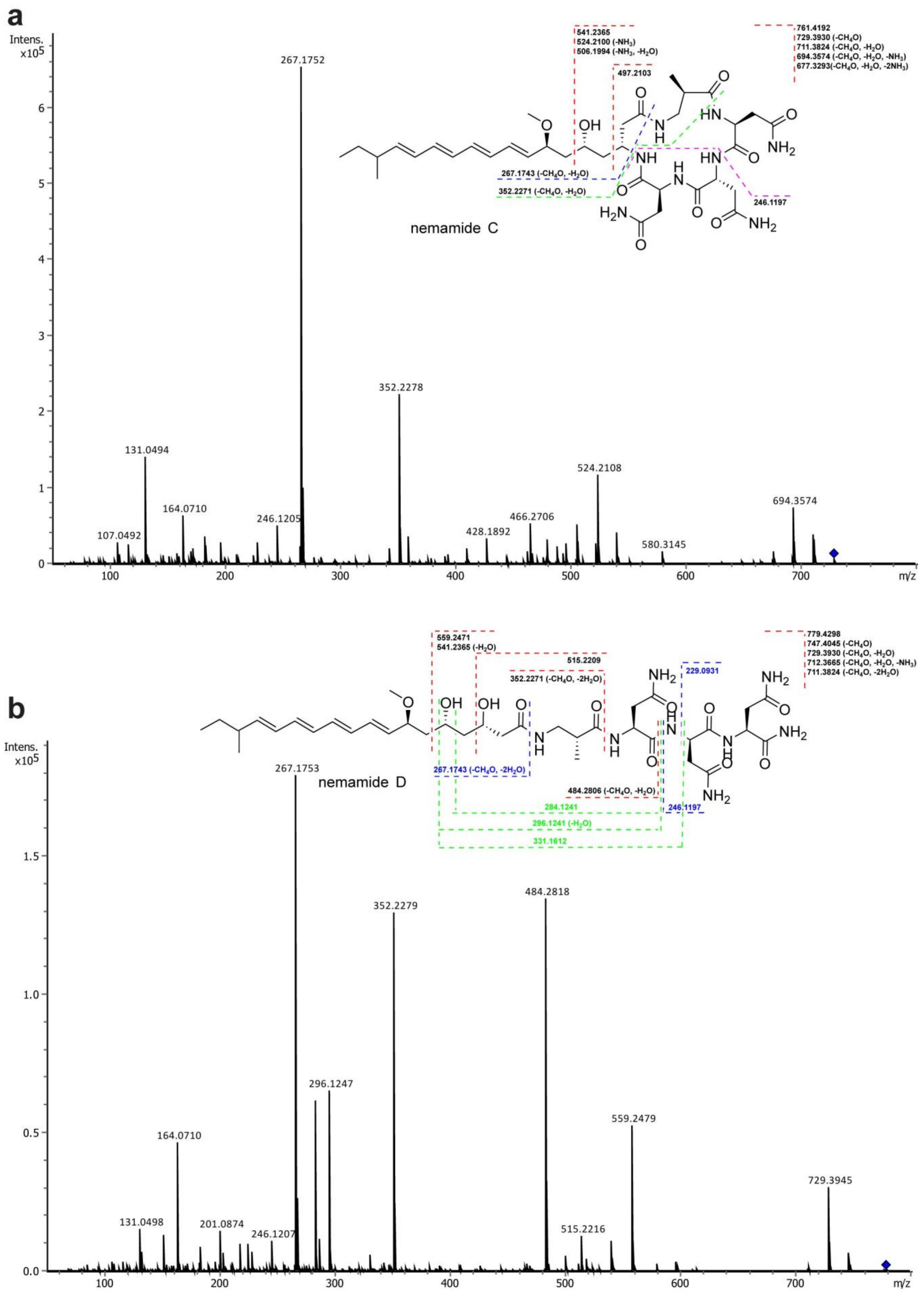
a
Intens. x10^5
541.2365
524.2100 (-NH3)
506.1994 (-NH3, -H2O)
497.2103
761.4192
729.3930 (-CH4O)
711.3824 (-CH4O, -H2O)
694.3574 (-CH4O, -H2O, -NH3)
677.3293(-CH4O, -H2O, -2NH3)
267.1743 (-CH4O, -H2O)
352.2271 (-CH4O, -H2O)
246.1197
nemamide C
267.1752
352.2278
131.0494
164.0710
107.0492
246.1205
428.1892
466.2706
524.2108
580.3145
694.3574
m/z
b
559.2471
541.2365 (-H2O)
515.2209
352.2271 (-CH4O, -2H2O)
229.0931
779.4298
747.4045 (-CH4O)
729.3930 (-CH4O, -H2O)
712.3665 (-CH4O, -H2O, -NH3)
711.3824 (-CH4O, -2H2O)
267.1743 (-CH4O, -2H2O)
484.2806 (-CH4O, -H2O)
284.1241
246.1197
296.1241 (-H2O)
331.1612
nemamide D
Intens. x10^5
267.1753
352.2279
484.2818
296.1247
164.0710
559.2479
729.3945
131.0498
201.0874
246.1207
515.2216
m/z

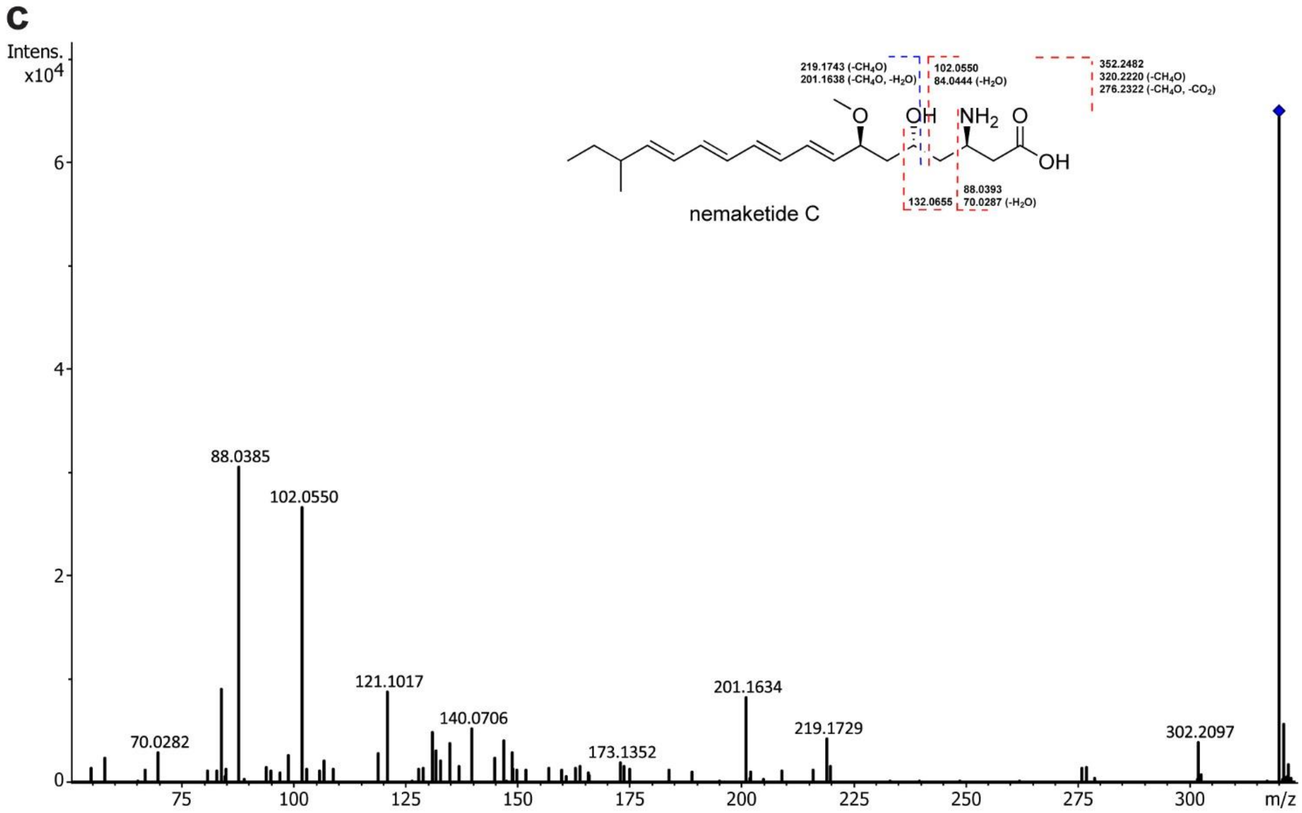


**Figure S1. MS/MS spectra of the nemamides of *P. pacificus*.** (**a**) MS/MS spectrum of nemamide C. (**b**) MS/MS spectrum of nemamide D. (**c**) MS/MS spectrum of nemaketide C.

**Figure S2. NMR spectra of nemamide C in dimethyl sulfoxide-$d_6$.** (**a**) $^1$H-NMR spectrum. (**b**) COSY spectrum. (**c**) dqf-COSY spectrum. (**d**) HSQC spectrum. (**e**) HMBC spectrum. (**f**) HMBC spectrum for the 170-185 ppm region.

**a**

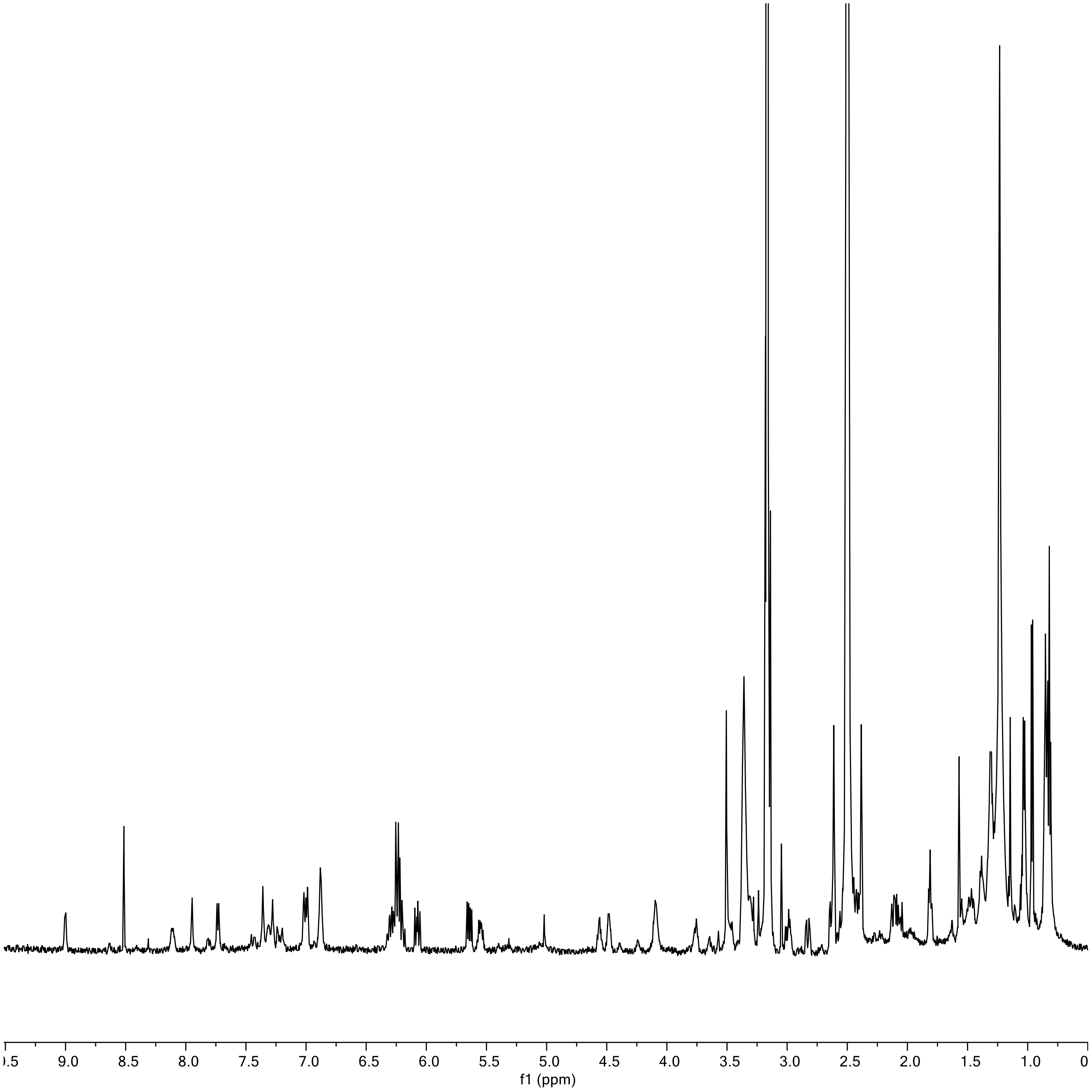

**b**

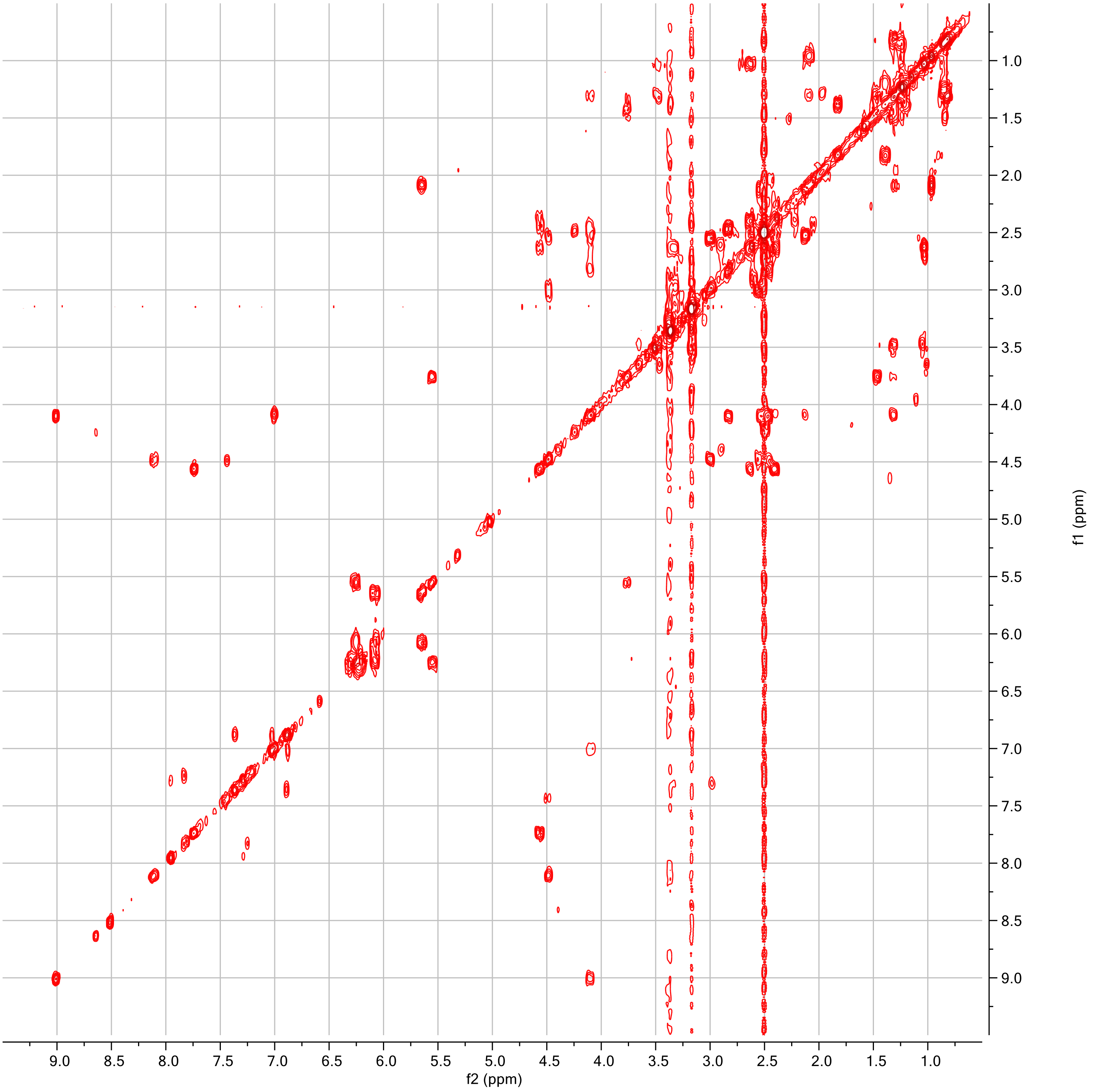

c

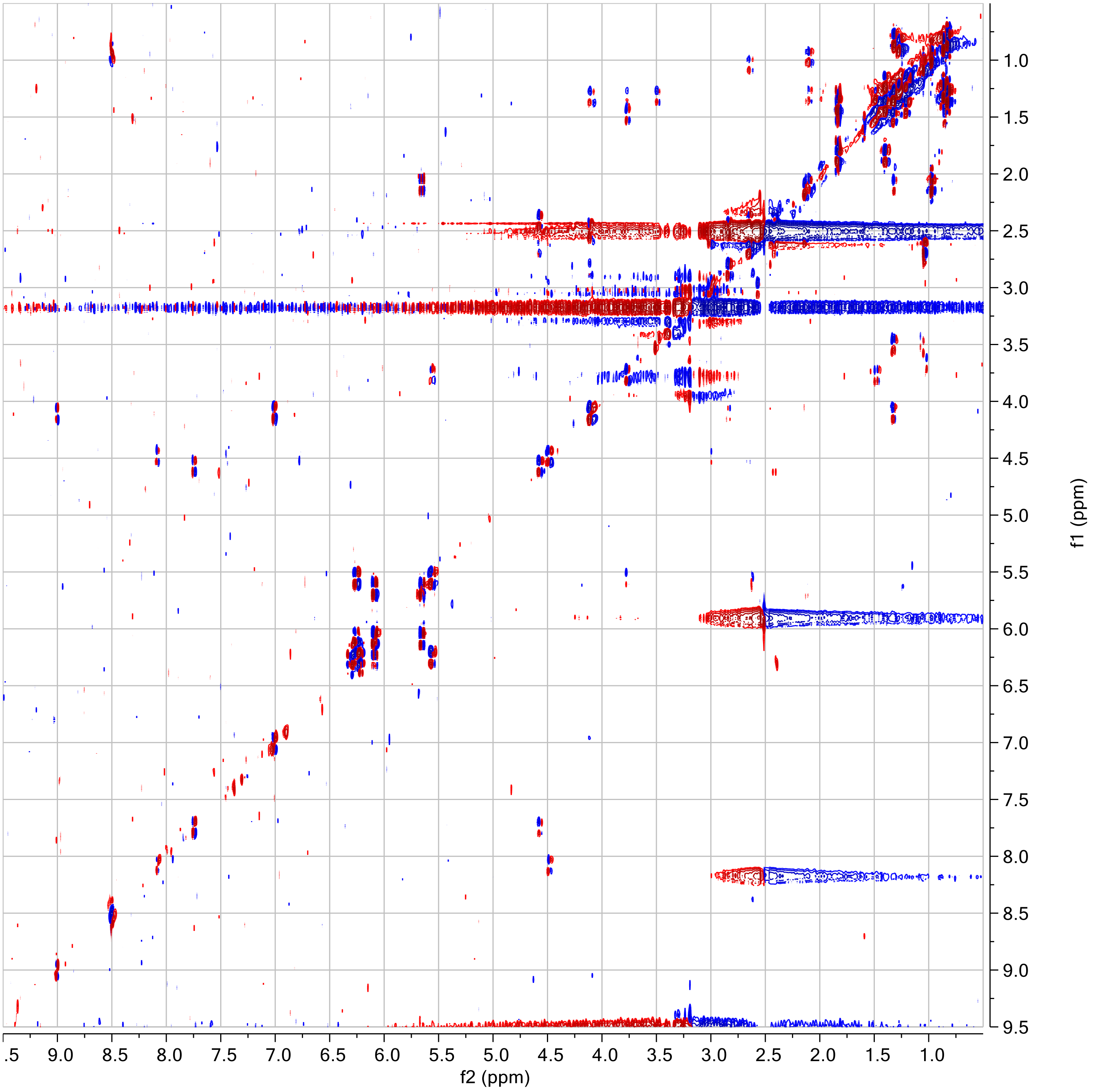

d

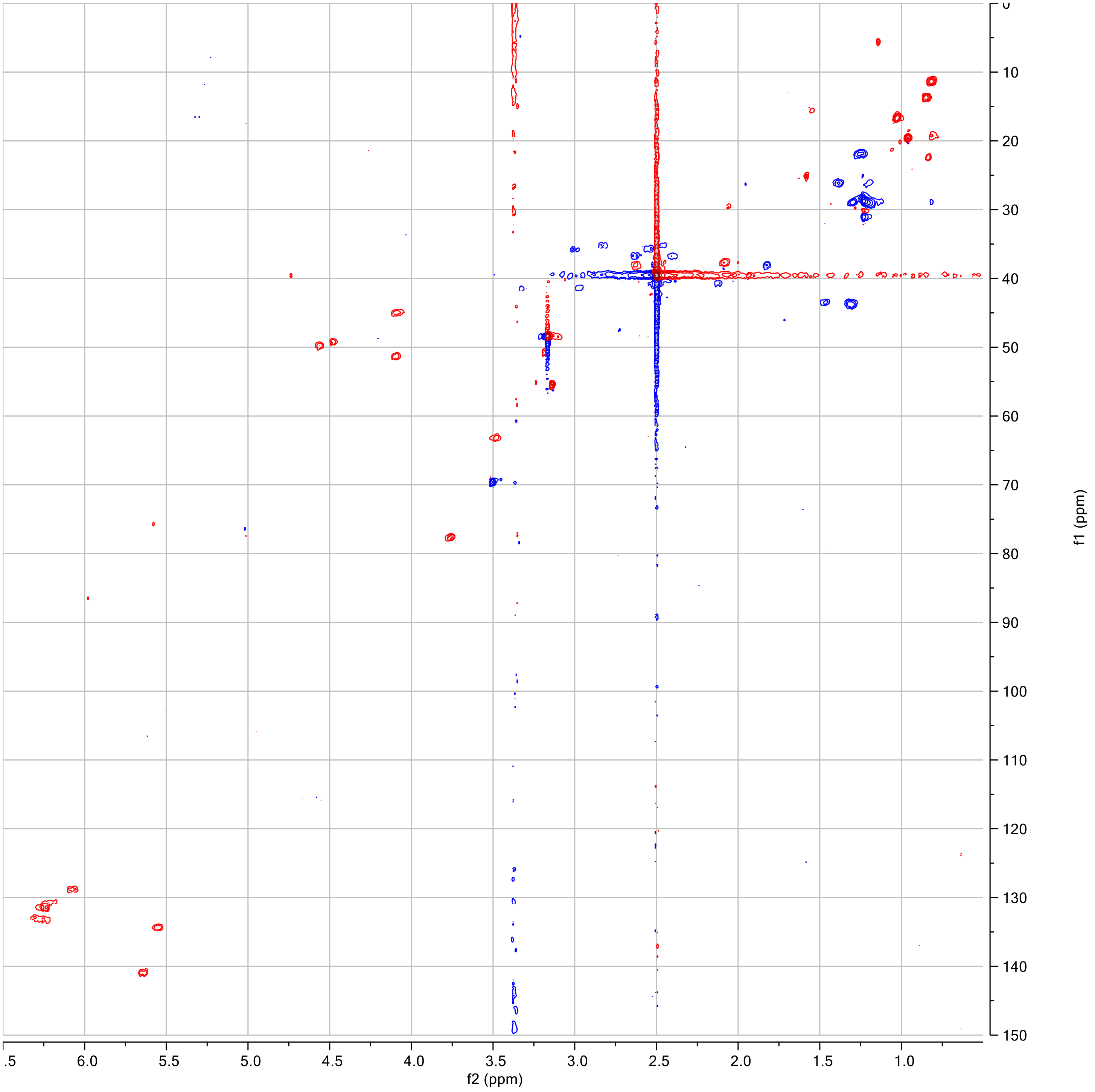

**e**

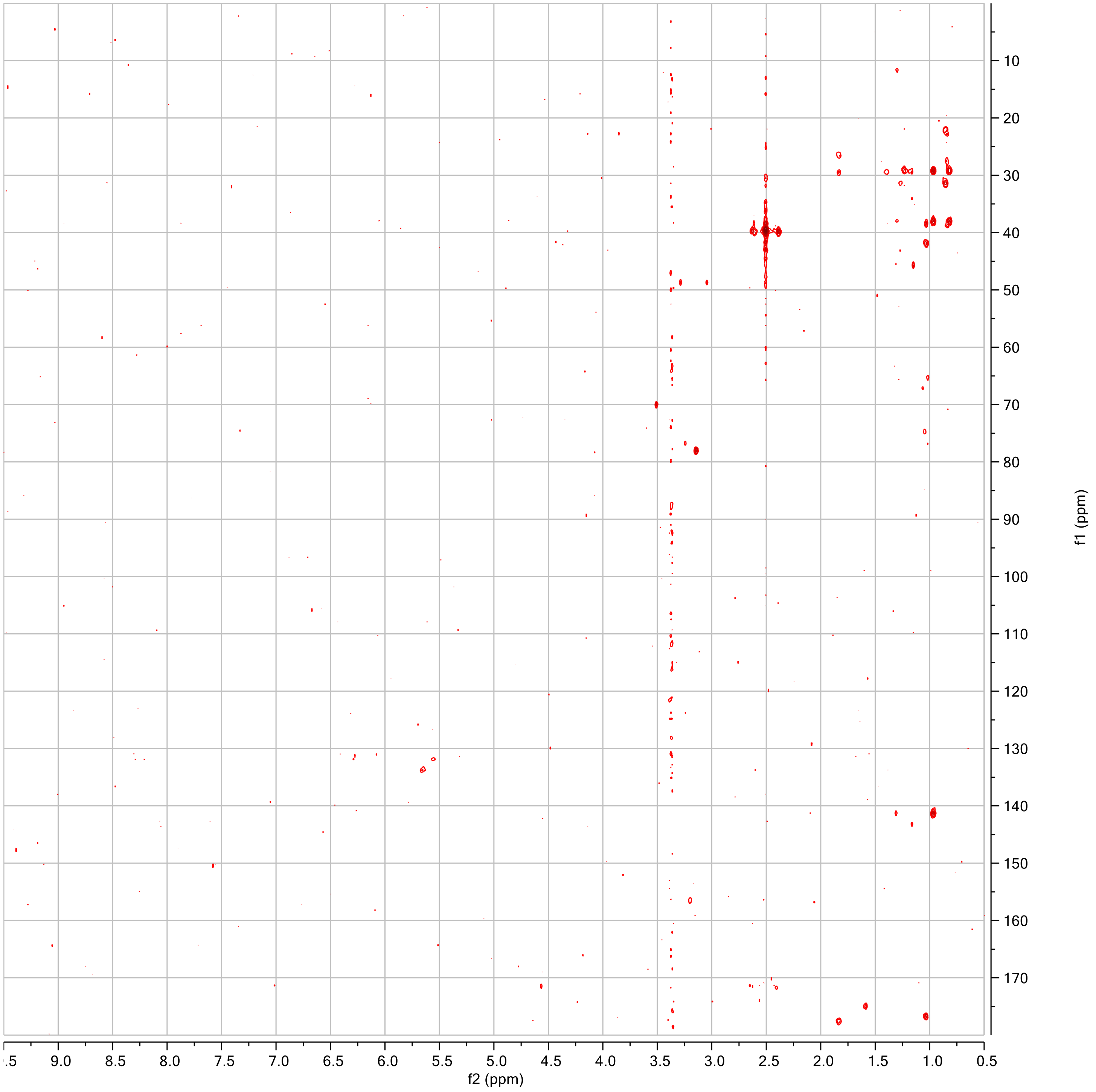

**f**

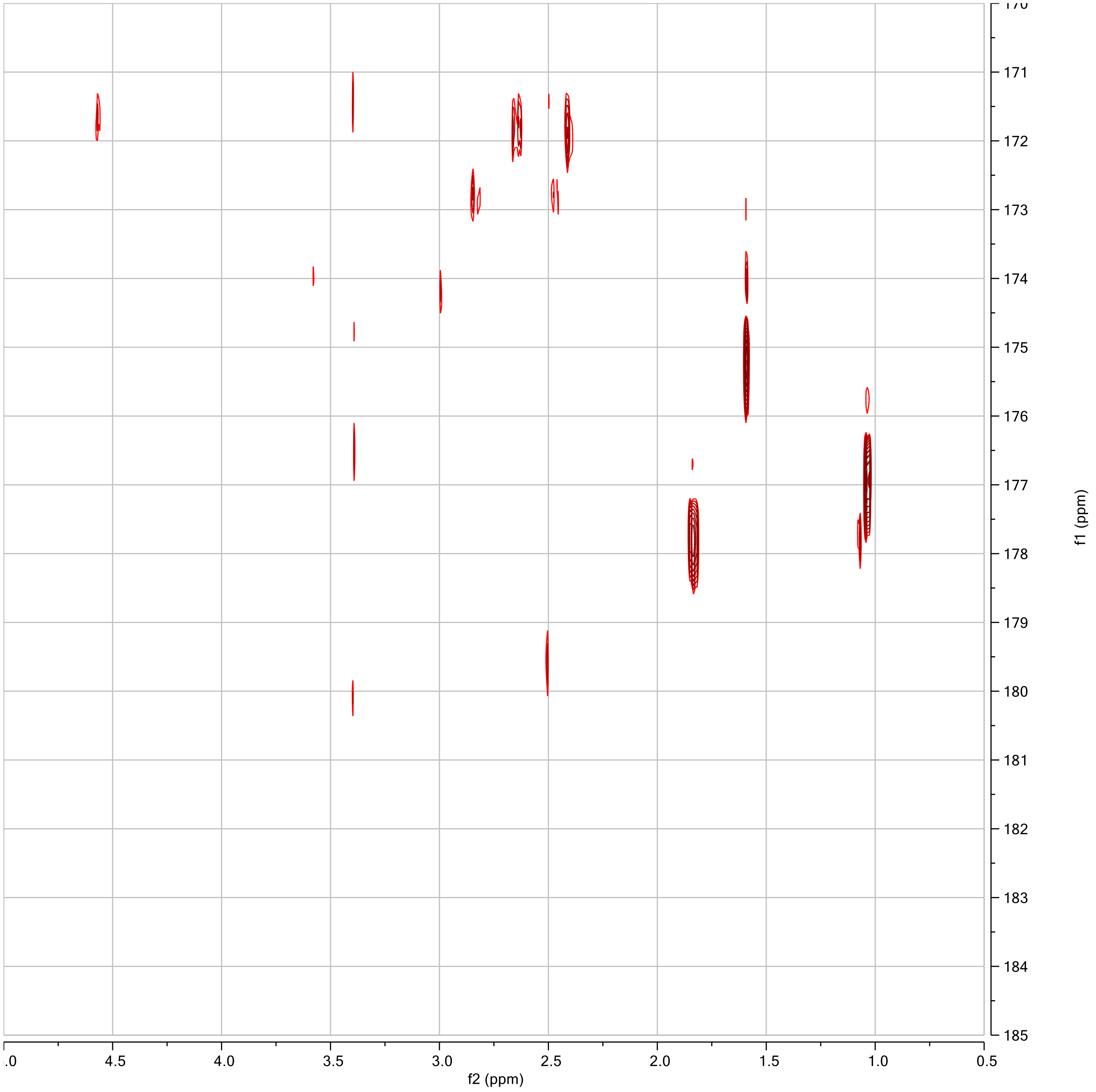

**Table S1. $^{1}H$ and $^{13}C$ NMR data derived from $^{1}H$, dqf-COSY, HSQC, and HMBC spectra for nemamide C in dimethyl sulfoxide-$d_6$.**

| # | $\delta_H$ ($J$ (Hz)) | $\delta_C$ | HMBC |
|---|---|---|---|
| 1 | | 171.4 | |
| 2 | 4.56, td ($J_{2,3a}$= 8.9; $J_{2,3b}$= 4.0) | 49.8 | $C_1$ |
| 2-NH | 7.76, d ($J_{2,2\text{-NH}}$= 8.0) | | $C_5$ |
| 3a | 2.41, m | 36.7 | $C_1$, $C_2$, $C_4$ |
| 3b | 2.63, m | 36.7 | |
| 4 | | 171.6 | |
| 4-NH2a* | N.D. | | |
| 4-NH2b* | N.D. | | |
| 5 | | 171.4 | |
| 6 | 4.48, m | 49.3 | $C_5$ |
| 6-NH | 8.11 br.s ($J_{6,6\text{-NH}}$= 7.8) | | |
| 7a | 2.55, overlap | 35.7 | |
| 7b | 3.00, m | | $C_5$, $C_8$ |
| 8 | | 174.4 | |
| 8-NH2a* | N.D. | | |
| 8-NH2b* | N.D. | | |
| 9 | | N.D. | |
| 10 | 4.10, overlap | 51.6 | |
| 10-NH | 9.00, d ($J_{10,10\text{-NH}}$= 6.0) | | |
| 11a | 2.47, overlap | 35.2 | $C_{10}$, $C_{12}$ |
| 11b | 2.83, dd ($J_{11a,11b}$ = 16.0, $J_{10,11b}$ = 4.5) | 35.2 | $C_{12}$ |
| 12 | | 172.5 | |
| 12-NH2a* | N.D. | | |
| 12-NH2b* | N.D. | | |
| 13 | | 176.8 | |
| 14 | 2.63, overlap | 38.7 | |
| 15a | 2.98, overlap | 42.3 | |
| 15b | 3.31, overlap | 42.3 | |
| 15-NH | 7.31, overlap | | |
| 16 | 1.04, d ($J_{14,16}$= 7.1) | 17.3 | $C_{13}$, $C_{14}$, $C_{15}$ |
| 17 | | N.D. | |
| 18a | 2.12, m ($J_{18a,18b}$ = 12.6, $J_{18a,19}$ = 3.9) | 40.7 | |
| 18b | 2.56, m | 40.7 | |
| 19 | 4.08, overlap | 45.4 | |
| 19-NH | 7.02, d ($J_{19,19\text{-NH}}$= 8.8) | | |
| 20 | 1.32, overlap | 44.7 | $C_{21}$ |
| 21 | 3.50, m ($J_{21,22a}$= 7.6) | 63.8 | |
| 21-OH | 3.65, br.s | | |
| 22a | 1.32, overlap | 44.1 | |
| 22b | 1.47, m ($J_{22b,23}$= 7.6) | 44.1 | |
| 23 | 3.75, td ($J_{23,25}$= 7.6) | 78.2 | |
| 24 | 3.14, s | 56.2 | $C_{23}$ |
| 25 | 5.56, m ($J_{25,26}$= 14.0) | 135.0 | $C_{26}$ |
| 26 | 6.25, overlap | 131.4 | |
| 27 | 6.22-6.29, overlap | 130.7-132.9, overlap | |
| 28 | 6.22-6.29, overlap | 130.7-132.9, overlap | |
| 29 | 6.22-6.29, overlap | 130.7-132.9, overlap | |
| 30 | 6.25, overlap | 133.1 | |
| 31 | 6.09, dd ($J_{30,31}$= 10.1) | 129.5 | |
| 32 | 5.65, dd ($J_{31,32}$= 15.1) | 140.9 | $C_{30}$ |
| 33 | 2.09, overlap ($J_{32,33}$= 8.1) | 38.3 | $C_{31}$, $C_{32}$ |
| 34 | 1.31, overlap | 29.6 | $C_{33}$, $C_{35}$, $C_{36}$ |
| 35 | 0.83, t ($J_{34,35}$= 7.4) | 12.1 | $C_{33}$, $C_{34}$ |
| 36 | 0.97, d ($J_{33,36}$= 6.7) | 20.3 | $C_{32}$, $C_{33}$, $C_{34}$ |

* Although amide protons were observed in the spectrum, the absence of HMBC correlations prevented them from being assigned to specific asparagines in nemamide C.

**Table S2. Comparison of $^1$H NMR chemical shifts of the polyketide chain of nemamide C with the polyketide chains of nemamide A and of euglenatides D and E.**

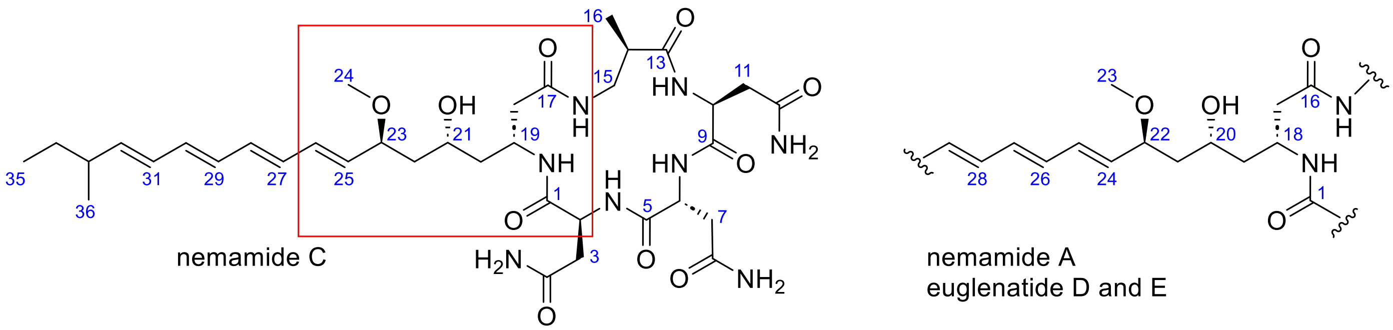


| # | **nemamide A**[1] $\delta_H$, mult., ($J$= Hz) | **euglenatide D**[2] $\delta_H$, mult., ($J$= Hz) | **euglenatide E**[2] $\delta_H$, mult., ($J$= Hz) | # | **nemamide C** $\delta_H$, mult., ($J$= Hz) |
|---|---|---|---|---|---|
| 17a | 2.18, br.d ($J_{17a,17b}$= 14.5) | 2.16, dd ($J_{17a,17b}$ = 12.8, $J_{17a,18}$ = 3.1) | 2.16, dd ($J_{17a,17b}$ = 13.8, $J_{17a,18}$ = 3.3) | 18a | 2.12, m ($J_{18a,18b}$ = 12.6, $J_{18a,19}$ = 3.9) |
| 17b | 2.47, overlap ($J_{17b,18}$= 9.8) | 2.47, overlap ($J_{17b,18}$= 9.6) | 2.47, overlap | 18b | 2.56, m |
| 18 | 4.08, m ($J_{18,19}$= 6.7) | 4.08, m ($J_{18,19}$= 6.5) | 4.08, m ($J_{18,19}$= 6.5) | 19 | 4.08, overlap |
| 18-NH | 7.05, brs ($J_{18,18\text{-}NH}$= 8.2) | 6.92, overlap | 6.92, overlap | 19-NH | 7.02, d ($J_{19,19\text{-}NH}$= 8.8) |
| 19 | 1.32, m ($J_{19,20}$ = 6.7) | 1.32, m | 1.32, m | 20 | 1.32, overlap |
| 20 | 3.51, m ($J_{20,21a}$= 8.2) | 3.51, m | 3.51, m | 21 | 3.50, m ($J_{21,22a}$= 7.6) |
| 21a | 1.30, m ($J_{21a,21b}$=14.5) | 1.30, m | 1.30, m | 22a | 1.32, overlap |
| 21b | 1.48, m ($J_{21b,22}$= 9.8) | 1.49, m | 1.48, m | 22b | 1.47, m ($J_{22b,23}$= 7.6) |
| 22 | 3.76, m ($J_{22,24}$= 7.5) | 3.75, m | 3.75, m | 23 | 3.75, td ($J_{23,25}$= 7.6) |
| 23 | 3.13, s | 3.13, s | 3.13, s | 24 | 3.14, s |
| 24 | 5.51, dd ($J_{24,25}$= 15.4) | 5.51, dd ($J_{24,25}$= 14.7, $J_{22,24}$= 7.7) | 5.51, dd ($J_{24,25}$= 14.6, $J_{22,24}$= 7.6) | 25 | 5.56, m ($J_{25,26}$= 14.0) |
| 25 | 6.19, dd ($J_{25,26}$= 11.2) | 6.20, dd ($J_{25,26}$= 10.5) | 6.19, dd ($J_{25,26}$= 10.2) | 26 | 6.25, m |

**Table S3. Comparison of $^{1}$H NMR chemical shifts of the peptide head of nemamide C with the peptide head of nemamide A and euglenatides B, C, and E.**

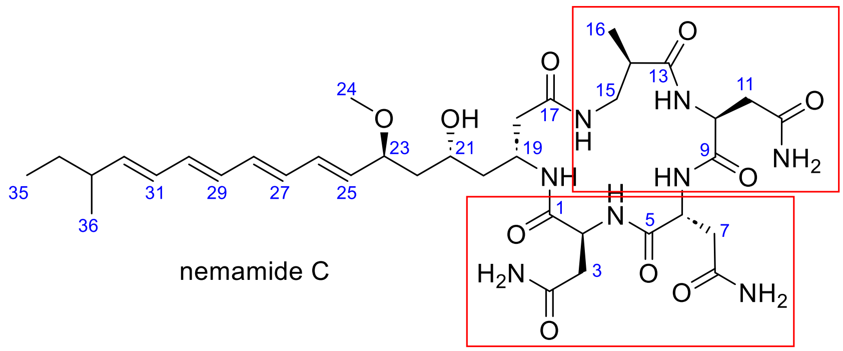

nemamide C

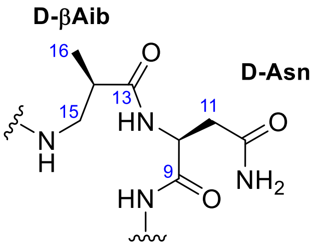


euglenatide B, C, and E

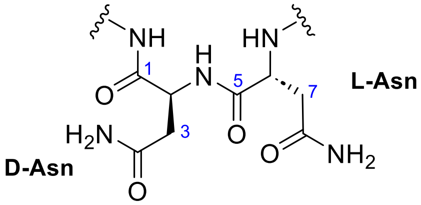


nemamide A

| # | nemamide A[1] $\delta_H$, mult., ($J$= Hz) | # | euglenatide B[2] $\delta_H$, mult., ($J$= Hz) | euglenatide C[2] $\delta_H$, mult., ($J$= Hz) | euglenatide E[2] $\delta_H$, mult., ($J$= Hz) | nemamide C $\delta_H$, mult., ($J$= Hz) |
|---|---|---|---|---|---|---|
| 2 | 4.51, m ($J_{2,3a}$= 9.0, $J_{2,3b}$= 6.9) | 2 | 4.32, m | 4.32 m | 4.30 m | 4.56, td ($J_{2,3a}$= 8.9; $J_{2,3b}$= 4.0) |
| 2-NH | 7.46, br.d ($J_{2,2\text{-}NH}$= 8.1) | 2-NH | 7.65, d ($J_{2,2\text{-}NH}$= 7.8) | 7.56, d ($J_{2,2\text{-}NH}$= 7.8) | 7.64, d ($J_{2,2\text{-}NH}$= 7.8) | 7.76 d ($J_{2,2\text{-}NH}$= 8.0) |
| 3a | 2.45, overlap ($J_{3a,3b}$= 15.9) | 3a | 1.71, m | 1.71 m | 1.69 m | 2.41, m |
| 3b | 2.61, overlap | 3b | 1.86, m | 1.86 m | 1.87 m | 2.63, m |
| 6 | 4.45, m ($J_{6,7a}$= 4.9; $J_{6,7b}$= 6.6) | 6 | 4.45, m | 4.45 m | 4.42 m | 4.48, m |
| 6-NH | 8.42, br.s ($J_{6,6\text{-}NH}$ = 8.2) | 6-NH | 7.77, d ($J_{6,6\text{-}NH}$ = 7.9) | 7.76, d ($J_{6,6\text{-}NH}$ = 8.5) | 7.87, d ($J_{6,6\text{-}NH}$ = 7.9) | 8.11 br.s ($J_{6,6\text{-}NH}$= 7.8) |
| 7a | 2.56, overlap ($J_{7a,7b}$ = 16.6) | 7a | 2.57, dd ($J_{7a,7b}$ = 16.9, $J_{6,7a}$ = 3.7) | 2.57, dd ($J_{7a,7b}$ = 17.0, $J_{6,7a}$ = 3.6) | 2.57, dd ($J_{7a,7b}$ = 16.9, $J_{6,7a}$ = 3.7) | 2.55, overlap |
| 7b | 2.95, dd | 7b | 2.94, dd ($J_{7a,7b}$ = 16.8, $J_{6,7b}$ = 5.0) | 2.94, dd ($J_{7a,7b}$ = 16.9, $J_{6,7b}$ = 5.0) | 3.01, dd ($J_{7a,7b}$ = 16.8, $J_{6,7b}$ = 5.0) | 3.00, m |
| 10 | 4.22, m ($J_{10,11a}$= 6.7; $J_{10,11b}$= 9.6) | 10 | 4.06, m | 4.06, m | 4.11, m | 4.10, overlap |
| 10-NH | 8.86, br.s ($J_{10,10\text{-}NH}$= 3.4) | 10-NH | 9.02, d ($J_{10,10\text{-}NH}$= 6.5) | 9.02, d ($J_{10,10\text{-}NH}$= 6.3) | 8.96, d ($J_{10,10\text{-}NH}$= 6.5) | 9.00, d ($J_{10,10\text{-}NH}$= 6.0) |
| 11a | 2.44, overlap ($J_{11a,11b}$ = 17.8) | 11a | 2.47, dd ($J_{11a,11b}$ = 15.9, $J_{10,11a}$ = 4.7) | 2.47, dd ($J_{11a,11b}$ = 15.5, $J_{10,11a}$ = 4.0) | 2.47, dd ($J_{11a,11b}$ = 15.9, $J_{10,11a}$ = 4.7) | 2.47, overlap |
| 11b | 2.53, overlap | 11b | 2.86, dd ($J_{11a,11b}$ = 15.7, $J_{10,11b}$ = 4.0) | 2.86, dd ($J_{11a,11b}$ = 15.4, $J_{10,11b}$ = 3.9) | 2.87, dd ($J_{11a,11b}$ = 15.7, $J_{10,11b}$ = 4.0) | 2.83, dd ($J_{11a,11b}$ = 16.0, $J_{10,11b}$= 4.5) |
| 14a | 2.41, overlap ($J_{14a,15a}$ = 8.9) | 14 | 2.66, m | 2.66, m | 2.64, m | 2.63, overlap |
| 14b | 2.57, overlap ($J_{14a,14b}$ = 18.5) | 15a | 3.03, br.d ($J_{15a,15b}$ = 13.0) | 3.03, br.d ($J_{15a,15b}$ = 11.2) | 2.93, br.d ($J_{15a,15b}$ = 13.0) | 2.98, overlap |
| 15a | 3.15, overlap ($J_{15a,15b}$ = 15.9) | 15b | 3.27, m | 3.27, m | 3.26, m | 3.31, overlap |
| 15b | 3.42, overlap ($J_{14b,15b}$ = 9.6) | 15-NH | 7.06, s | 7.06, s | 6.96, overlap | 7.31, overlap |
| 15-NH | 7.73, br.s | 16 | 1.04, d ($J_{14,16}$= 7.1) | 1.02, d ($J_{14,16}$= 6.2) | 1.02, d ($J_{14,16}$= 7.1) | 1.04, d ($J_{14,16}$= 7.1) |

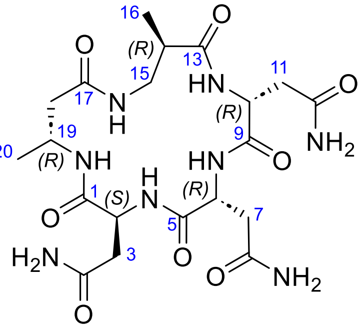

cyclic peptide **S1**

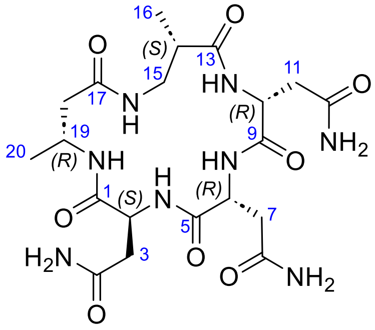

cyclic peptide **S2**

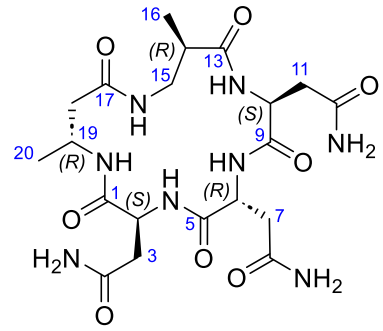

cyclic peptide **S3**

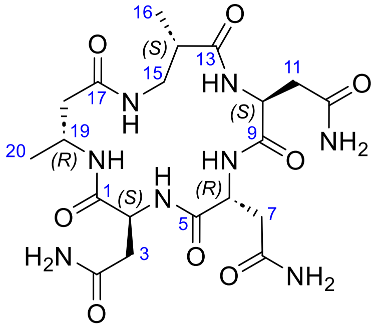

cyclic peptide **S4**

**Figure S3. Structures of the four model cyclic peptides (S1-S4).**

**Table S4. $^{1}H$ and $^{13}C$ NMR chemical shifts derived from $^{1}H$, DQF-COSY, HSQC, and HMBC spectra for the four cyclic peptides S1-S4 in dimethyl sulfoxide-$d_6$.**

| # | Cyclic Peptide **S1** (2*S*, 6*R*, 10*R*, 14*R*,19*R*) | | Cyclic Peptide **S2** (2*S*, 6*R*, 10*R*, 14*S*, 19*R*) | | Cyclic Peptide **S3** (2*S*, 6*R*, 10*S*, 14*R*, 19*R*) | | Cyclic Peptide **S4** (2*S*, 6*R*, 10*S*, 14*S*, 19*R*) | |
|---|---|---|---|---|---|---|---|---|
| | $\delta_H$ (*J* (Hz)) | $\delta_C$ | $\delta_H$ (*J* (Hz)) | $\delta_C$ | $\delta_H$ (*J* (Hz)) | $\delta_C$ | $\delta_H$ (*J* (Hz)) | $\delta_C$ |
| 1 | | 170.15 | | 170.40 | | 169.98 | | 169.90 |
| 2 | 4.44, m | 50.80 | 4.53, m | 50.32 | 4.51, m | 50.37 | 4.55, m | 50.09 |
| 2-NH | 7.43, brd, ($J_{2,2\text{-NH}}$ = 8.4) | | 7.58, brd, ($J_{2,2\text{-NH}}$ = 8.4) | | 7.75, brd, ($J_{2,2\text{-NH}}$ = 9.0) | | 8.21, brd, ($J_{2,2\text{-NH}}$ = 7.8) | |
| 3a | 2.46, dd, ($J_{3a,3b}$ = 15.6; $J_{2,3a}$ =9.6) | 37.05 | 2.43, overlap | 36.92 | 2.39, overlap | 37.19 | 2.38, dd, ($J_{3a,3b}$ = 16.2; $J_{2,3a}$ =6.6) | 35.93 |
| 3b | 2.60, dd, ($J_{3a,3b}$ = 16.2; $J_{2,3b}$ =3.6) | 37.05 | 2.66, dd, ($J_{3a,3b}$ = 15.6; $J_{2,3b}$ =4.2) | 36.92 | 2.64, dd, ($J_{3a,3b}$ = 15.6; $J_{2,3b}$ =4.8) | 37.19 | 2.67, overlap | 35.93 |
| 4 | | 171.58 | | 171.75 | | 171.80 | | 172.20 |
| 4-$NH_2$a | 6.88, brs | | 6.81, brs | | 6.90, brs | | 6.98, brs | |
| 4-$NH_2$b | 7.21, brs | | 7.18, brs | | 7.35, brs | | 7.40, brs | |
| 5 | | 170.67 | | 170.51 | | 171.15 | | 170.76 |
| 6 | 4.33, m | 50.00 | 4.38, m | 49.41 | 4.43, m | 49.57 | 4.48, m | 49.65 |
| 6-NH | 8.37, brd, ($J_{6,6\text{-NH}}$ = 9.0) | | 8.12, brd, ($J_{6,6\text{-NH}}$ = 8.4) | | 7.93, brd, ($J_{6,6\text{-NH}}$ =9.0) | | 7.88, brd, ($J_{6,6\text{-NH}}$ = 9.6) | |
| 7a | 2.64, dd, ($J_{7a,7b}$ = 16.2; $J_{6,7a}$ =4.2) | 35.93 | 2.57, dd, ($J_{7a,7b}$ = 15.6; $J_{6,7a}$ =4.2) | 35.88 | 2.58, overlap | 36.00 | 2.51, overlap | 36.05 |
| 7b | 2.86, dd, ($J_{7a,7b}$ = 16.8; $J_{6,7b}$ =6.0) | 35.93 | 2.98, dd, ($J_{7a,7b}$ = 16.8; $J_{6,7b}$ =4.8) | 35.88 | 3.00, dd, ($J_{7a,7b}$ = 17.4; $J_{6,7b}$ =4.8) | 36.00 | 3.08, dd, ($J_{7a,7b}$ = 17.4; $J_{6,7b}$ =4.2) | 36.05 |
| 8 | | 173.92 | | 174.21 | | 174.30 | | 174.20 |
| 8-$NH_2$a | 7.29, brs | | 7.01, brs | | 7.42, brs | | 6.88, brs | |
| 8-$NH_2$b | 7.78, brs | | 7.45, brs | | 7.90, brs | | 7.40, brs | |
| 9 | | 171.45 | | 171.84 | | 170.03 | | 170.71 |
| 10 | 4.28, m | 52.30 | 4.18, m | 53.23 | 4.13, m | 51.87 | 4.43, m | 49.07 |
| 10-NH | 8.54, brd, ($J_{10,10\text{-NH}}$ = 4.8) | | 8.23, brd, ($J_{10,10\text{-NH}}$ = 4.8) | | 8.93, brd, ($J_{10,10\text{-NH}}$ = 6.6) | | 8.21, brd, ($J_{10,10\text{-NH}}$ = 7.8) | |
| 11a | 2.49, overlap | 36.21 | 2.50, overlap | 35.77 | 2.50, overlap | 35.56 | 2.48, overlap | 35.82 |
| 11b | 2.49, overlap | 36.21 | 2.56, dd, ($J_{11a,11b}$ = 15.6; $J_{10,11b}$ =4.2) | 35.77 | 2.82, dd, ($J_{11a,11b}$ = 15.6; $J_{10,11b}$ =4.2) | 35.56 | 2.68, overlap | 35.82 |
| 12 | | 171.02 | | 171.49 | | 172.50 | | 173.00 |
| 12-$NH_2$a | 6.99, brs | | 7.40, brs | | 6.89, brs | | 7.48, brs | |
| 12-$NH_2$b | 7.44, brs | | 7.86, brs | | 7.02, brs | | 7.92, brs | |
| 13 | | 176.53 | | 177.61 | | 176.45 | | 176.46 |
| 14 | 2.70, m | 38.38 | 2.52, m | 42.53 | 2.61, m | 38.79 | 2.55, m | 41.78 |
| 15a | 3.12, m | 42.39 | 3.23, m | 42.25 | 3.06, m | 41.84 | 3.00, m | 42.08 |
| 15b | 3.25, m | 42.39 | 3.36, m | 42.25 | 3.35, overlap | 41.84 | 3.40, m | 42.08 |
| 15-NH | 6.94, t, ($J_{15,15\text{-NH}}$ = 4.2) | | 7.45, m | | 6.96, m | | 7.40, m | |
| 16 | 1.01, d, ($J_{14,16}$ = 7.2) | 16.62 | 1.07, d, ($J_{14,16}$ = 7.2) | 16.17 | 1.02, d, ($J_{14,16}$ = 7.2) | 16.78 | 1.00, d, ($J_{14,16}$ = 7.2) | 16.87 |
| 17 | | 170.58 | | 170.85 | | 170.88 | | 170.85 |
| 18a | 2.20, dd, ($J_{18a,18b}$ = 12.6; $J_{19,18a}$ =3.0) | 42.32 | 2.16, dd, ($J_{18a,18b}$ = 13.2; $J_{19,18a}$ =2.4) | 42.09 | 2.09, dd, ($J_{18a,18b}$ = 13.2; $J_{19,18a}$ =3.6) | 42.56 | 2.07, dd, ($J_{18a,18b}$ = 13.8; $J_{19,18a}$ =2.4) | 42.04 |
| 18b | 2.35, m | 42.32 | 2.46, m | 42.09 | 2.50, overlap | 42.56 | 2.58, m | 42.04 |
| 19 | 3.95, m | 43.54 | 3.93, m | 43.88 | 3.97, m | 43.70 | 3.98, m | 43.87 |

**Table S5. Differences between the cyclic peptides S1-S4 and nemamide C in terms of $^{1}H$ and $^{13}C$ NMR chemical shifts.***

| # | $\delta_{H(peptide)}$- $\delta_{H(nemamideC)}$ | | | | $\delta_{C(peptide)}$- $\delta_{C(nemamideC)}$ | | | |
|---|---|---|---|---|---|---|---|---|
| | Cyclic Peptide **S1** (2*S*,6*R*, 10*R*,14*R*, 19*R*) | Cyclic Peptide **S2** (2*S*,6*R*, 10*R*,14*S*, 19*R*) | Cyclic Peptide **S3** (2*S*,6*R*, 10*S*,14*R*, 19*R*) | Cyclic Peptide **S4** (2*S*,6*R*, 10*S*,14*S*, 19*R*) | Cyclic Peptide **S1** (2*S*,6*R*, 10*R*,14*R*, 19*R*) | Cyclic Peptide **S2** (2*S*,6*R*, 10*R*,14*S*, 19*R*) | Cyclic Peptide **S3** (2*S*,6*R*, 10*S*,14*R*, 19*R*) | Cyclic Peptide **S4** (2*S*,6*R*, 10*S*,14*S*, 19*R*) |
| 1 | | | | | -1.41 | -1.16 | -1.58 | -1.66 |
| 2 | -0.12 | -0.03 | -0.05 | -0.01 | 1.03 | 0.23 | 0.60 | 0.32 |
| 3a | 0.02 | -0.01 | 0.05 | -0.06 | 0.29 | 0.16 | 0.43 | -0.83 |
| 3b | -0.03 | 0.03 | 0.01 | 0.04 | 0.29 | 0.16 | 0.43 | -0.83 |
| 4 | | | | | -0.15 | 0.02 | 0.07 | 0.47 |
| 5 | | | | | -0.70 | 0.86 | -0.22 | 0.61 |
| 6 | -0.15 | -0.10 | -0.05 | 0.00 | 0.75 | 0.16 | 0.32 | 0.40 |
| 7a | 0.10 | 0.03 | 0.04 | -0.03 | 0.23 | 0.18 | 0.30 | 0.35 |
| 7b | -0.14 | -0.02 | 0.00 | 0.08 | 0.23 | 0.18 | 0.30 | 0.35 |
| 8 | | | | | -0.25 | 0.04 | 0.13 | 0.03 |
| 9 | | | | | | | | |
| 10 | 0.19 | 0.10 | 0.04 | 0.34 | 0.9 | 1.83 | 0.47 | -2.33 |
| 11a | 0.02 | 0.03 | 0.03 | 0.01 | 1.01 | 0.57 | 0.36 | 0.62 |
| 11b | -0.33 | -0.26 | 0.00 | -0.14 | 1.01 | 0.57 | 0.36 | 0.62 |
| 12 | | | | | 1.56 | 1.09 | -0.08 | 0.42 |
| 13 | | | | | -0.41 | 0.67 | -0.49 | -0.48 |
| 14 | 0.08 | -0.10 | -0.01 | -0.07 | 0.36 | 3.83 | 0.77 | 3.08 |
| 15a | 0.15 | 0.26 | 0.09 | 0.03 | 1.05 | 0.91 | 0.41 | 0.74 |
| 15b | -0.06 | 0.05 | 0.04 | 0.10 | 1.05 | 0.91 | 0.41 | 0.74 |
| 16 | -0.03 | 0.04 | -0.01 | -0.03 | 0.00 | -0.45 | 0.16 | 0.25 |
| 17 | | | | | | | | |
| 18a | 0.08 | 0.04 | -0.03 | -0.05 | 2.79 | 2.56 | 3.03 | 2.51 |
| 18b | -0.23 | -0.10 | 0.06 | 0.02 | 2.79 | 2.56 | 3.03 | 2.51 |
| 19 | -0.14 | -0.15 | -0.12 | -0.11 | -1.42 | -1.08 | -1.26 | -1.09 |
| 20 | -0.33 | -0.33 | -0.35 | -0.35 | -23.36 | -23.35 | -22.91 | -23.05 |

* $^{1}H$ and $^{13}C$ NMR chemical shifts of nemamide C were subtracted from the corresponding chemical shifts of the four cyclic peptides. If $\delta_{H(cyclic\ peptide)}$- $\delta_{H(nemamideC)} > 0.1$, the value is highlighted in red. If $\delta_{C(cyclic\ peptide)}$- $\delta_{C(nemamideC)} > 1$, the value is highlighted in red.

**Figure S4. NMR spectra of ascarene A in methanol-*d₄*.** (**a**) $^{1}$H-NMR spectrum of pure ascarene A. (**b**) $^{1}$H-NMR spectrum. (**c**) COSY spectrum. (**d**) dqf-COSY spectrum. (**e**) HSQC spectrum. (**f**) HMBC spectrum. (**g**) NOESY spectrum.

**a**

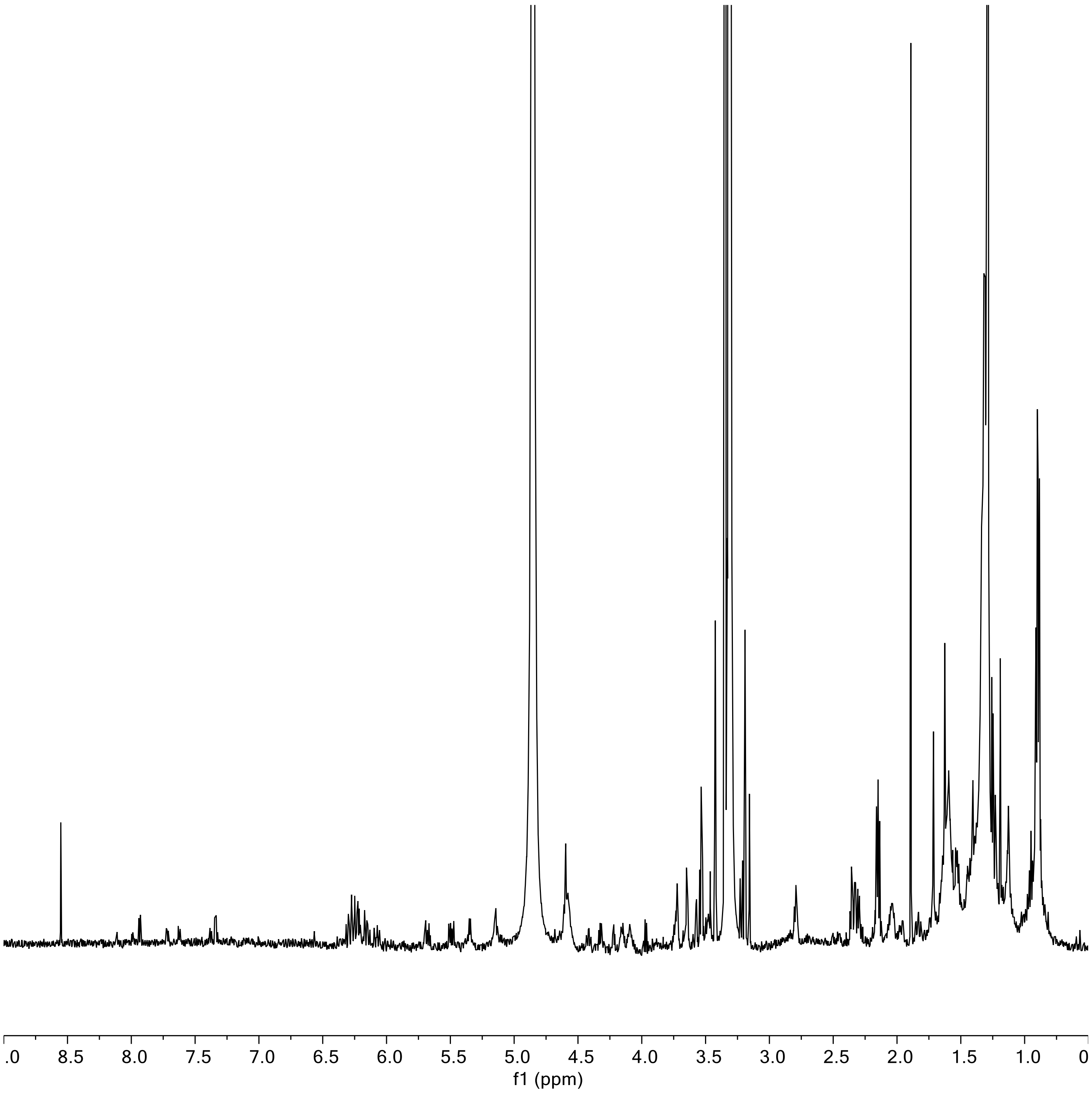

b

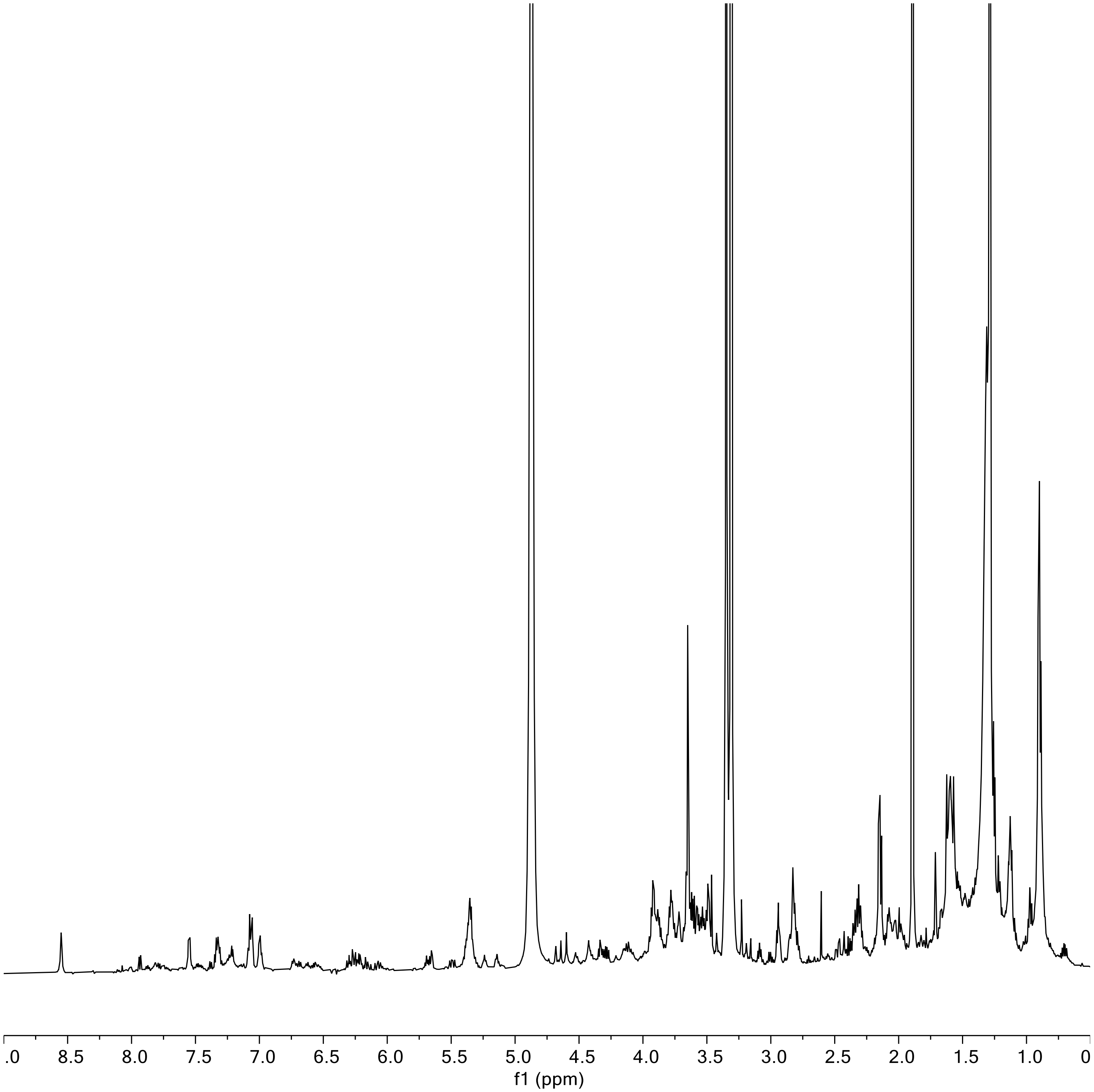

c

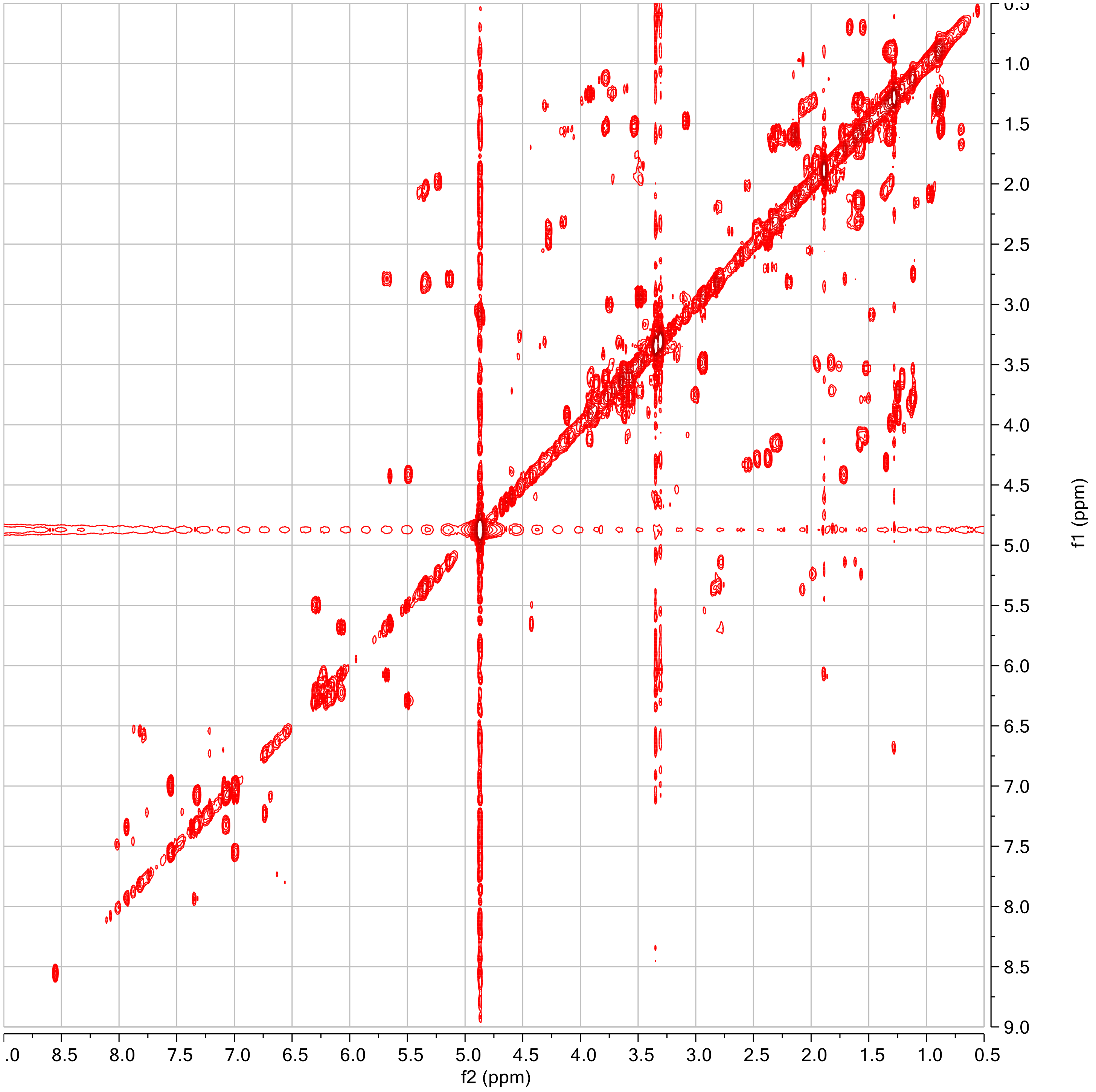

d

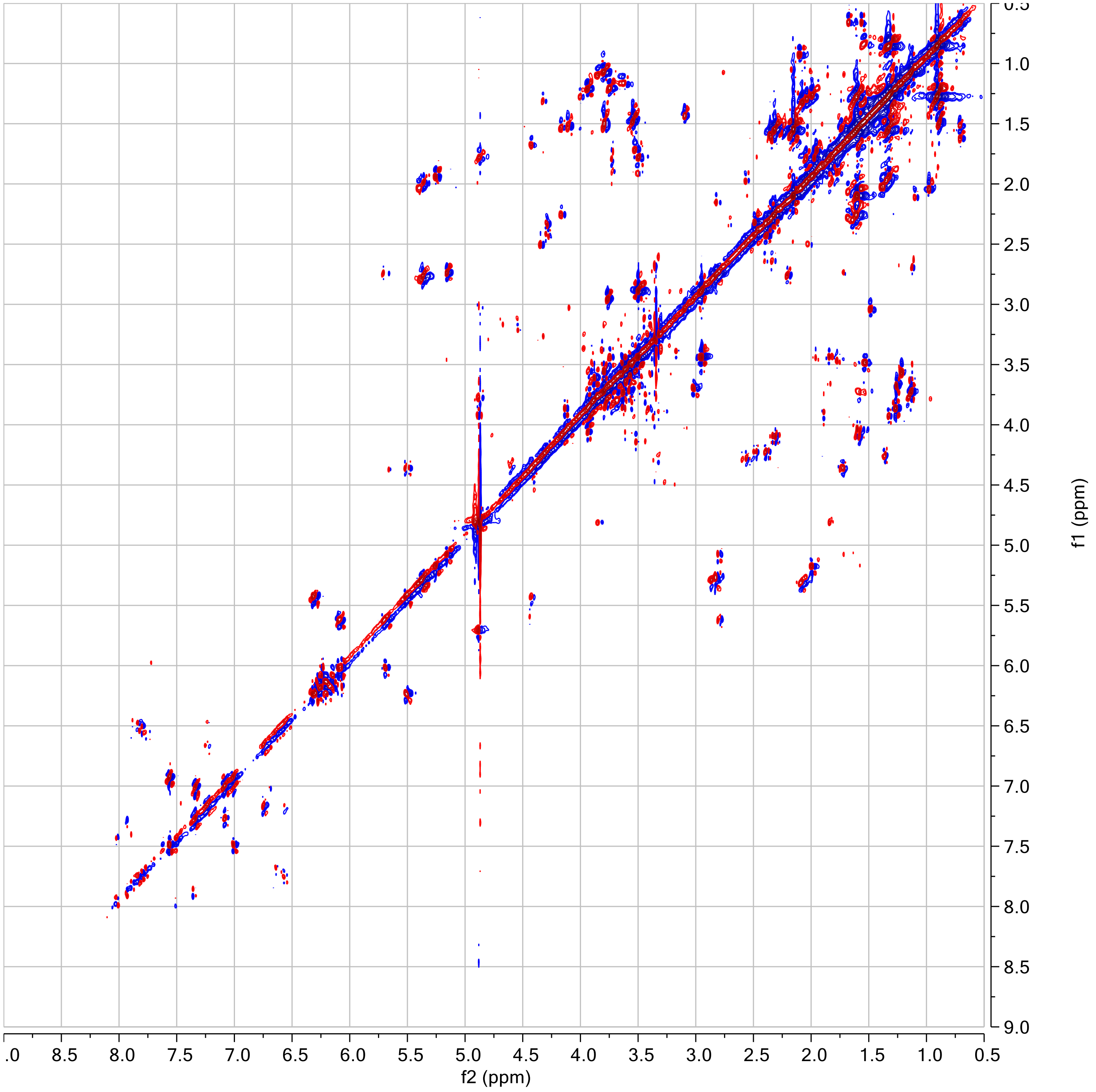

e

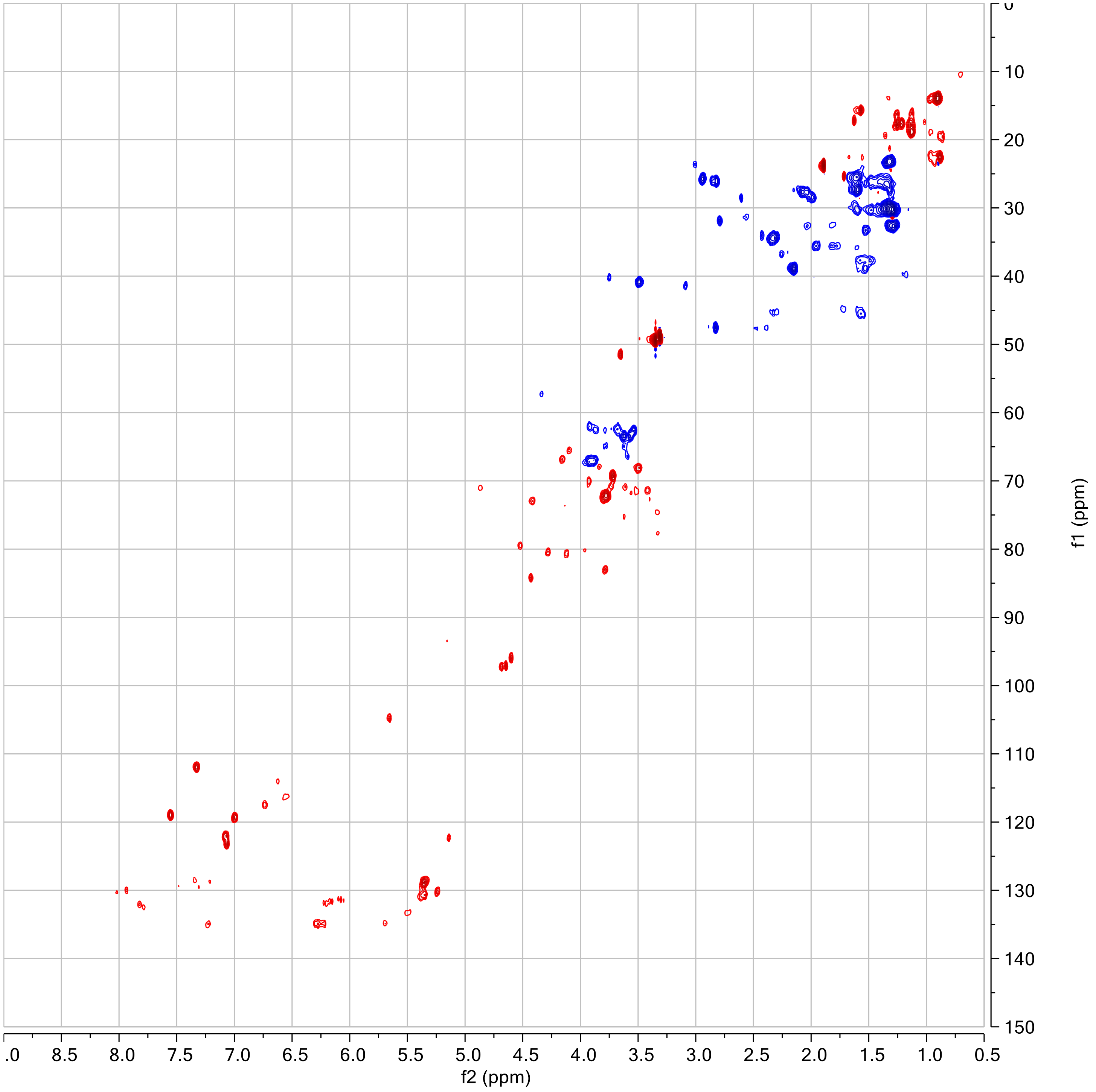

f2 (ppm)
f1 (ppm)
.0
8.5
8.0
7.5
7.0
6.5
6.0
5.5
5.0
4.5
4.0
3.5
3.0
2.5
2.0
1.5
1.0
0.5
0
10
20
30
40
50
60
70
80
90
100
110
120
130
140
150

**f**

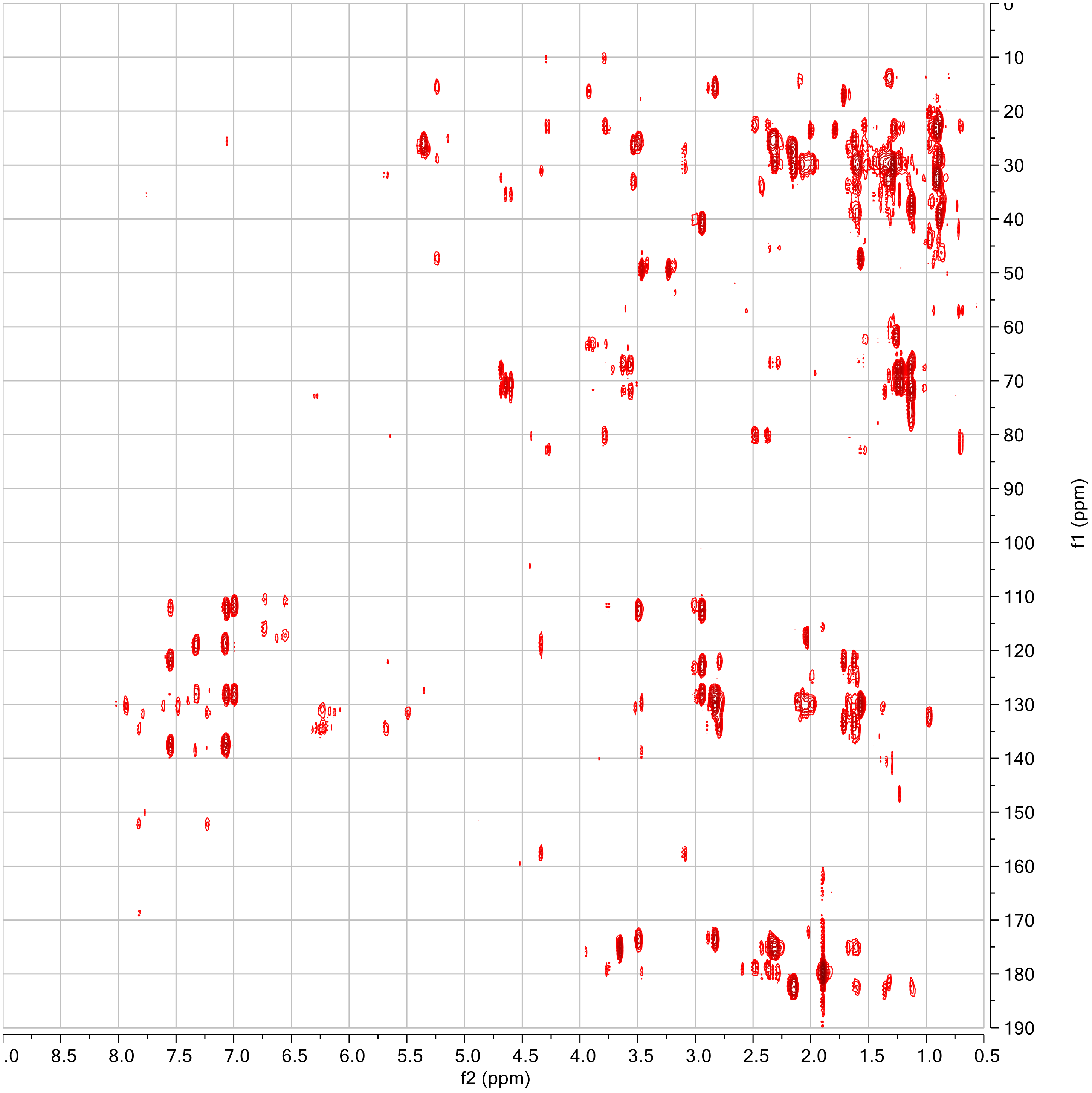

g

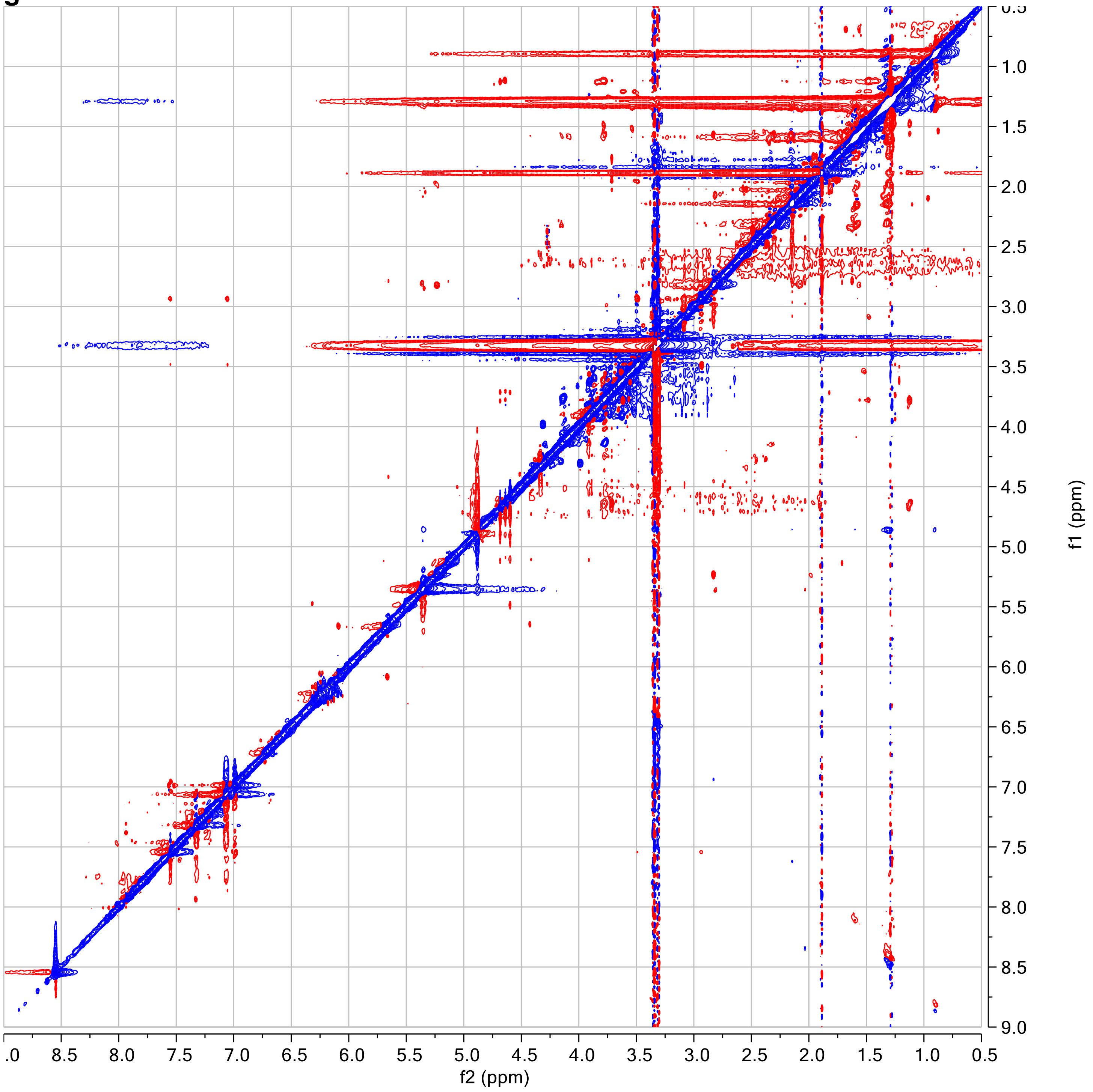

**Table S6.** $^{1}H$ **and** $^{13}C$ **NMR data derived from** $^{1}H$**, dqf-COSY, HSQC, and HMBC spectra for ascarene A in methanol-$d_4$.**

| # | $\delta_H$ ($J$ (Hz)) | $\delta_C$ | HMBC |
|---|---|---|---|
| 1 | | 180.1 | |
| 2a | 2.28, m | 45.4 | $C_1$, $C_3$ |
| 2b | 2.34, m | 45.4 | $C_1$, $C_3$ |
| 3 | 4.15, m | 66.8 | |
| 4a | 1.54, overlap | 45.7 | $C_3$, $C_5$ |
| 4b | 1.57, overlap | 45.7 | $C_3$, $C_5$ |
| 5 | 4.09, m | 65.5 | |
| 6a | 1.59, overlap | 44.8 | $C_5$ |
| 6b | 1.72, overlap | 44.8 | $C_5$ |
| 7 | 4.42, td ($J_{7,8}$ =9.4, $J_{7,6a/b}$ = 3.0) | 73.0 | |
| 8 | 5.49, dd ($J_{8,9}$ =15.1) | 133.3 | $C_{10}$ |
| 9 | 6.30, dd ($J_{9,10}$ = 10.4) | 135.0 | $C_{11}$ |
| 10 | 6.22, dd ($J_{10,11}$ =15.1) | 131.9 | $C_8$, $C_9$, $C_{11}$, $C_{12}$ |
| 11 | 6.27, dd ($J_{11,12}$ = 10.4) | 135.0 | $C_9$, $C_{12}$, $C_{13}$ |
| 12 | 6.16, dd ($J_{12,13}$ = 15.1) | 131.7 | $C_{14}$ |
| 13 | 6.23, dd ($J_{13,14}$ = 10.4) | 135.0 | $C_{11}$, $C_{12}$, $C_{14}$, $C_{15}$ |
| 14 | 6.08, dd like ($J_{14,15}$ = 15.1) | 131.4 | $C_{12}$ |
| 15 | 5.68, dt ($J_{15,16}$ = 7.0) | 134.8 | $C_{13}$, $C_{16}$, $C_{17}$ |
| 16 | 2.79, t like ($J_{16,17}$ = 7.0) | 31.9 | $C_{14}$, $C_{17}$, $C_{18}$ |
| 17 | 5.14, m | 122.3 | |
| 18 | | 133.4 | |
| 19 | 1.62, br. s | 17.3 | $C_{17}$, $C_{18}$, $C_{20}$ |
| 20 | 1.71, d ($J_{17,20}$ = 0.8) | 25.4 | $C_{17}$, $C_{18}$, $C_{19}$ |
| 1′ | 4.60, m | 96.0 | $C_7$, $C_{2'}$, $C_{3'}$, $C_{5'}$ |
| 2′ | 3.70, overlap | 69.4 | |
| 3′ax | 1.82, ddd ($J_{2',3'ax}$= 2.9, $J_{3'ax,4'}$= 11.3, $J_{3'ax,3'eq}$= 12.9) | 35.5 | |
| 3′eq | 1.97, ddd like ($J_{2',3'eq}$= 4.0) | 35.5 | |
| 4′ | 3.50, overlap | 68.2 | |
| 5′ | 3.73, overlap | 70.8 | |
| 6′ | 1.25, d ($J_{5',6'}$ = 6.2) | 17.7 | $C_{4'}$, $C_{5'}$ |

**a**

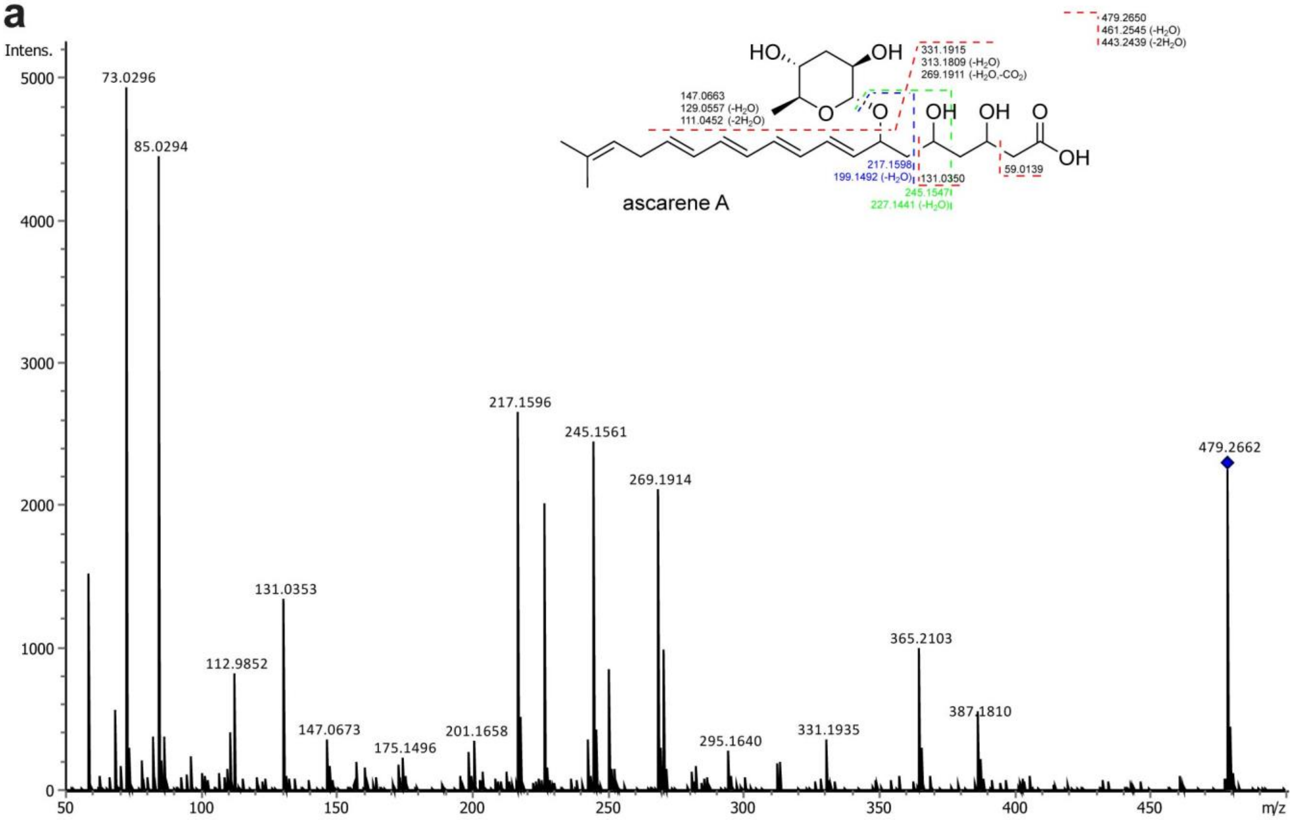
Intens.
5000
4000
3000
2000
1000
0
50
100
150
200
250
300
350
400
450
m/z
73.0296
85.0294
112.9852
131.0353
147.0673
175.1496
201.1658
217.1596
245.1561
269.1914
295.1640
331.1935
365.2103
387.1810
479.2662
ascarene A
HO
OH
O
O
OH
OH
O
OH
147.0663
129.0557 (-H2O)
111.0452 (-2H2O)
331.1915
313.1809 (-H2O)
269.1911 (-H2O,-CO2)
479.2650
461.2545 (-H2O)
443.2439 (-2H2O)
217.1598
199.1492 (-H2O)
131.0350
245.1547
227.1441 (-H2O)
59.0139

**b**

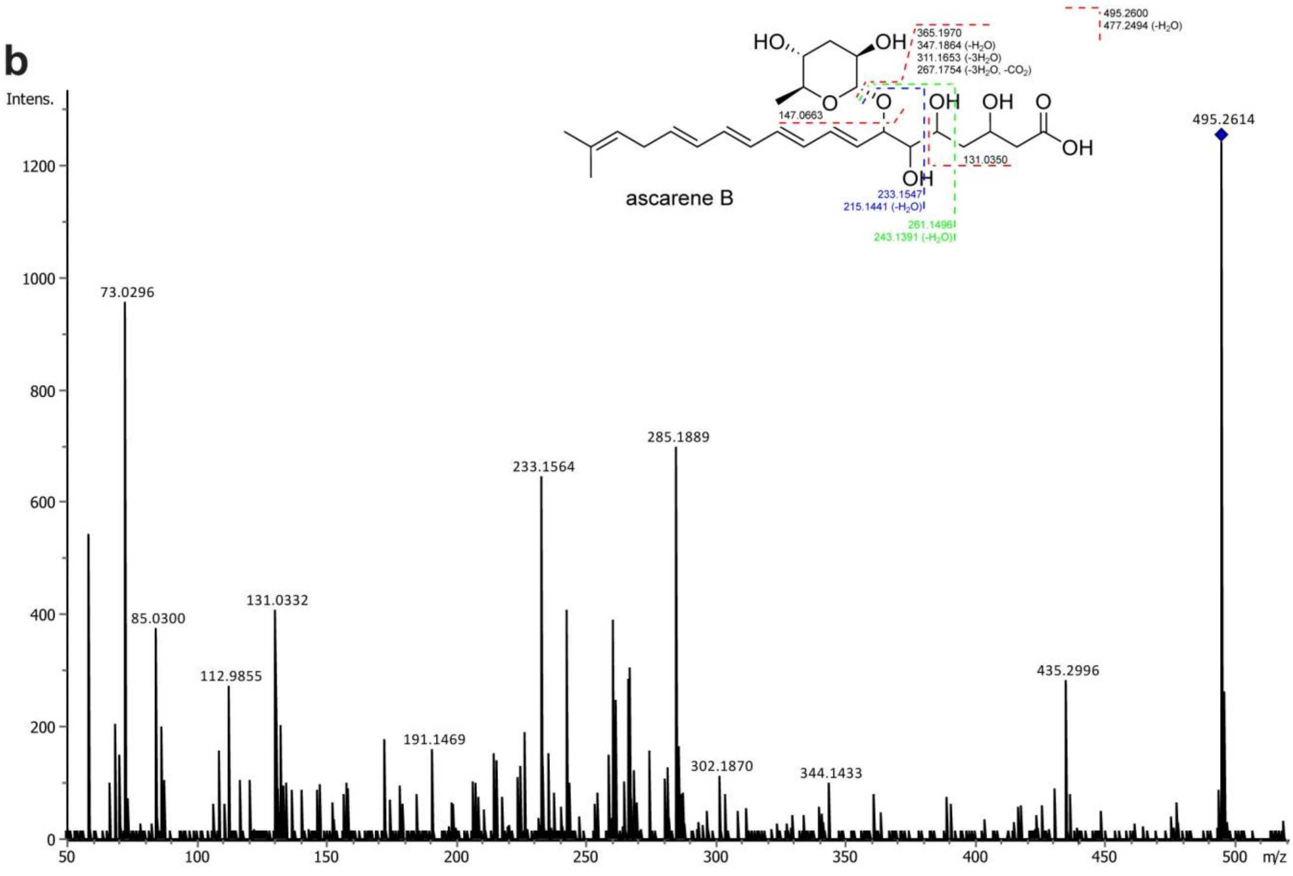
Intens.
1200
1000
800
600
400
200
0
50
100
150
200
250
300
350
400
450
500
m/z
73.0296
85.0300
112.9855
131.0332
191.1469
233.1564
285.1889
302.1870
344.1433
435.2996
495.2614
ascarene B
HO
OH
O
O
OH
OH
OH
O
OH
147.0663
365.1970
347.1864 (-H2O)
311.1653 (-3H2O)
267.1754 (-3H2O, -CO2)
495.2600
477.2494 (-H2O)
233.1547
215.1441 (-H2O)
131.0350
261.1496
243.1391 (-H2O)

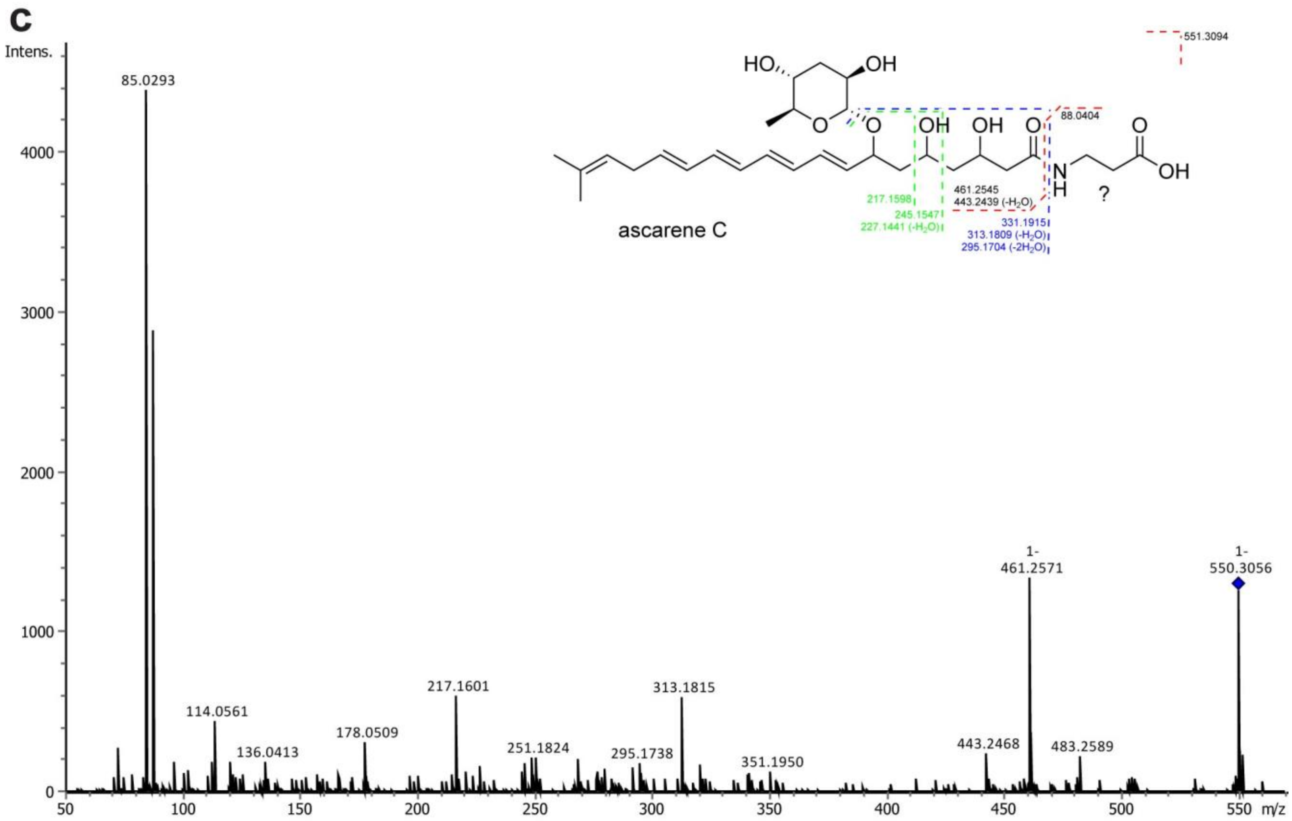


**Figure S5. MS/MS spectra of the ascarenes of *P. pacificus*.** MS/MS spectrum of ascarene A (**a**), ascarene B (**b**), and ascarene C (**c**).

**Table S7. Comparison of $^{1}$H and $^{13}$C NMR chemical shifts of the sugar of ascarene A with ascarylose of asc-C6-MK and paratose of part#9 in methanol-$d_4$.**

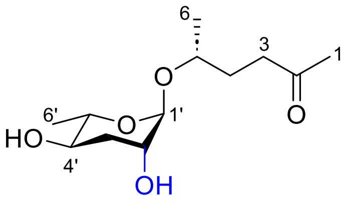


asc-C6-MK

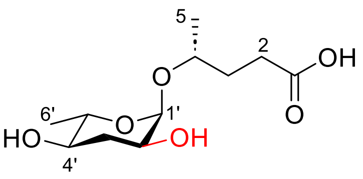


part#9

| | # | ascarene A | asc-C6-MK[3] | Δ | part#9[4] | Δ |
|---|---|---|---|---|---|---|
| $\delta_H$ | 1' | 4.60, m | 4.63 br s | -0.03 | 4.74, d ($J_{1',2'}$ = 3.9) | -0.14 |
| | 2' | 3.70, overlap | 3.69, dt ($J_{1',2'}$ = 1.6, $J_{2',3'ax}$= 3.6) | 0.01 | 3.60, ddd ($J_{2',3'ax}$= 12.1, $J_{2',3'eq}$= 5.5) | -0.10 |
| | 3'ax | 1.82, ddd ($J_{2',3'ax}$= 2.9, $J_{3'ax,4'}$= 11.3, $J_{3'ax,3'eq}$= 12.9) | 1.74, ddd ($J_{3'ax,4'}$= 11.2, $J_{3'ax,3'eq}$= 13.3) | 0.08 | 2.02 dt ($J_{2',3'ax}$= 10.9) | -0.20 |
| | 3'eq | 1.97, ddd like ($J_{2',3'eq}$= 4.0) | 1.94, dt ($J_{2',3'eq}$= 3.6) | 0.03 | 1.74, dt ($J_{3'eq,4'}$= 4.7, $J_{3'ax,3'eq}$= 12.3) | 0.23 |
| | 4' | 3.50, overlap | 3.51, ddd ($J_{3'eq,4}$= 3.6) | -0.01 | 3.15, ddd ($J_{4',5'}$= 9.4) | 0.35 |
| | 5' | 3.73, overlap | 3.55, dq ($J_{4',5'}$ = 9.3) | 0.18 | 3.58, dq ($J_{5',6'}$ = 6.1) | 0.15 |
| | 6' | 1.25, d ($J_{5',6'}$ = 6.2) | 1.21, d ($J_{5',6'}$ = 6.0) | 0.04 | 1.18 | 0.07 |
| $\delta_C$ | 1' | 96.0 | 97.4 | -1.4 | 95.9 | 0.1 |
| | 2' | 69.4 | 69.9 | -0.5 | 68.4 | 1.0 |
| | 3' | 35.5 | 36.0 | -0.5 | 36.8 | -1.3 |
| | 4' | 68.2 | 68.3 | -0.1 | 71.6 | -3.4 |
| | 5' | 70.8 | 71.3 | -0.5 | 69.9 | 0.9 |
| | 6' | 17.7 | 18.1 | -0.4 | 17.5 | 0.2 |

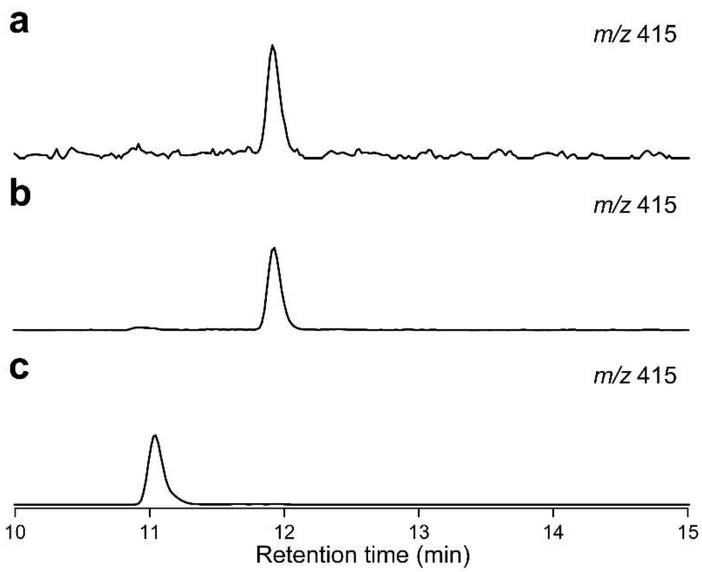


**Figure S6. Determination of absolute configuration of the sugar in ascarene A.** The EIC (*m/z* 415) of the derivatized sugar hydrolysate of ascarene A (**a**), the derivatized ascarylose standard (**b**), the derivatized tyvelose standard (**c**).

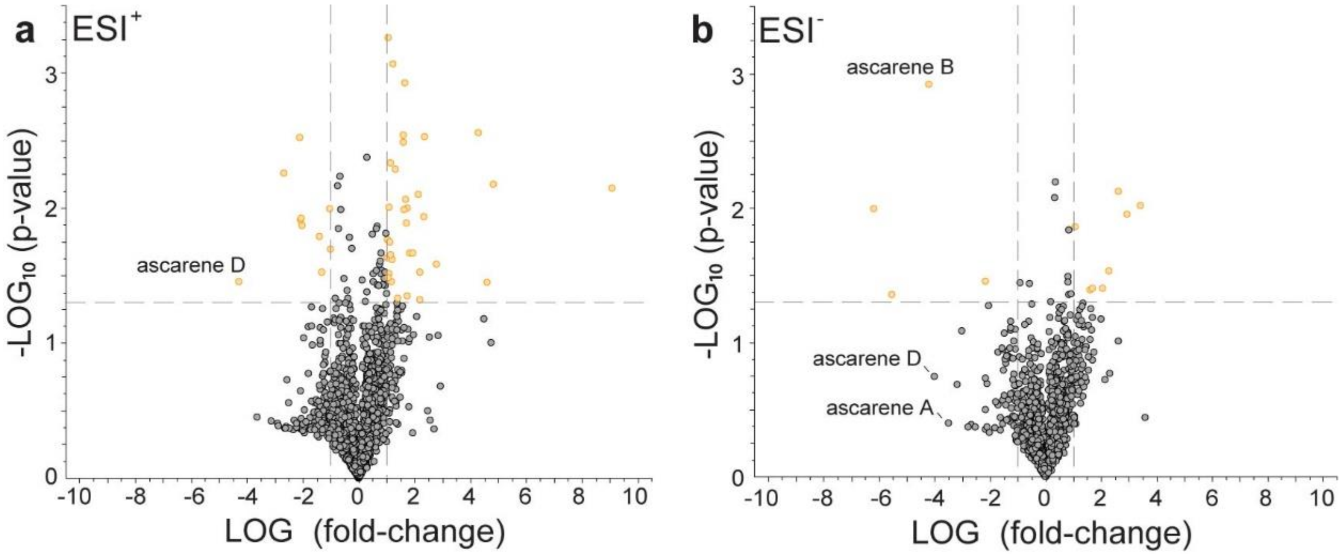


**Figure S7. Comparative metabolomics between wild-type and *pks-1(tu1297)* exometabolomes.** (**a**,**b**) Volcano plot comparing the average intensities of metabolites in wild-type versus *pks-1*(*tu1297*) exometabolomes in positive (ESI$^+$) (**a**) and negative (ESI$^-$) (**b**) modes. The x-axis indicates the log ratio of the area of a given peak in wild-type versus *pks-1*(*tu1297*), and the y-axis indicates the statistical significance. The ascarenes are indicated.

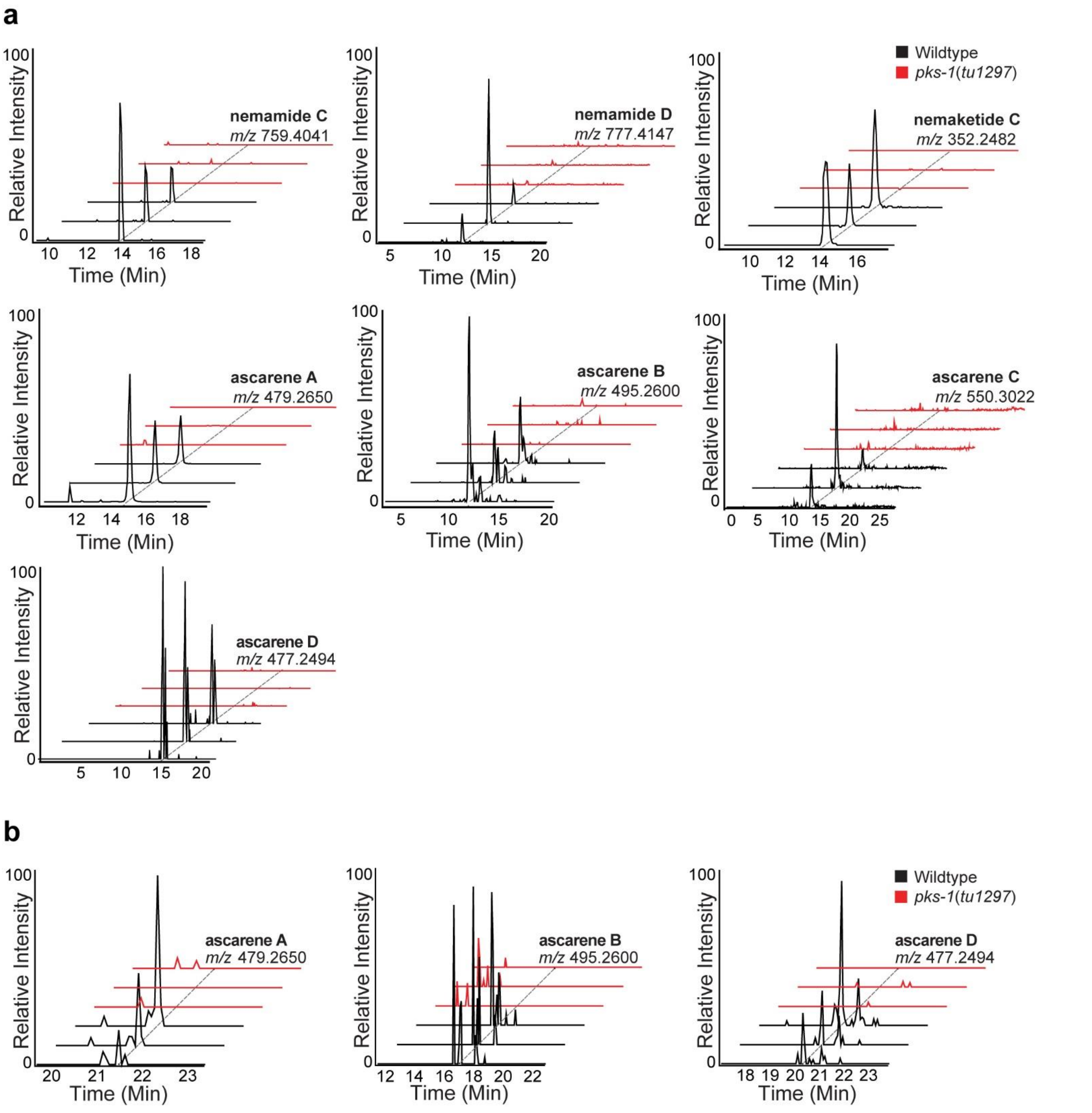


**Figure S8**. **Production of nemamides and ascarenes in the endometabolome and exometabolome of *P. pacificus*.** (**a**) Extracted ion chromatograms of nemamide C, nemamide D, nemaketide C, and ascarenes A-D in the endometabolome. (**b**) Extracted ion chromatograms of ascarenes A, B, and D in the exometabolome.
